# Morpho-electric and computational properties of three types of human hippocampal CA1 pyramidal neurons

**DOI:** 10.1101/2023.10.05.561029

**Authors:** Eline J. Mertens, Yoni Leibner, Jean Pie, Anna A. Galakhova, Femke Waleboer, Julia Meijer, Tim S. Heistek, René Wilbers, Djai Heyer, Natalia A. Goriounova, Sander Idema, Matthijs B. Verhoog, Brian E. Kalmbach, Brian R. Lee, Ryder P. Gwinn, Ed S. Lein, Eleonora Aronica, Jonathan Ting, Huibert D. Mansvelder, Idan Segev, Christiaan P.J. de Kock

## Abstract

Hippocampal pyramidal neuron activity underlies episodic memory and spatial navigation. Although extensively studied in rodents, extremely little is known about human hippocampal pyramidal neurons, even though human hippocampus underwent strong evolutionary reorganization and shows lower theta rhythm frequencies. To test whether biophysical and computational properties of human *CA1* pyramidal neurons can explain observed rhythms, we map the morpho-electric and computational properties of individual *CA1* pyramidal neurons in human, non-pathological hippocampal slices from neurosurgery. Human *CA1* pyramidal neurons have extensive dendrites and resonate at 2.9 Hz, optimally tuned to human theta frequencies. Morphological and biophysical properties reveal three cell types with distinct dendrite bifurcations and physiology. Data-driven biophysical models show that human *CA1* pyramidal neurons use i) computationally independent dendritic compartments, ii) preferential routing of electrical activity towards soma or dendritic tree and iii) non-linear input-output transformations. Across cell types, morpho-electric properties consistently increase computational richness in human *CA1* pyramidal neurons.

**Teaser:** Human *CA1* pyramidal neurons have large and intricate morphologies, which translates to complex computational properties.

## INTRODUCTION

Navigating through an environment and remembering the steps and events along the way rely on hippocampal function ^1,2^. Although the hippocampal complex and many of its basic functions are conserved across mammals ^3,4^, the human hippocampal complex in general and *CA1* cytoarchitecture specifically show a dramatic reorganization during evolution. The increase in size and altered subregional organization in the human hippocampus are the largest in primate evolution ^5,6^. These evolutionary changes are likely associated with the emergence of specialized human cognitive abilities, such as extraordinary cognitive flexibility ^7^. Not surprisingly, human cognitive decline is consistently linked to decreased function of the hippocampal complex ^8,9^.

Functionally, during spatial navigation and mnemonic processing, hippocampal networks generate a prominent theta rhythm ^10,11^ and phase-locking of single neurons to the theta rhythm is associated with human memory strength and spatial navigation capabilities ^12,13^. Impaired hippocampal theta rhythm directly affects spatial navigation and cognitive performance ^14,15^. Interestingly, the theta rhythm is the only oscillatory brain rhythm that inversely scales with brain size across mammals, in contrast to alpha, beta and gamma rhythms, which are similar across mammalian brains ^16,17^. In human hippocampus, theta rhythm is 1 – 4 Hz ^16–19^ compared to 4 – 10 Hz in rats ^2,20^. A slower theta rhythm directly impacts theories on human brain function, since the slower cycle could allow an increased number of neuronal assemblies to interact and lock to the same cycle ^16^. This could translate to the association of an increased number of items in working memory compared to rodents and thus increased human cognitive flexibility ^21,22^.

The ability of single neurons to respond with high selectivity to inputs at preferred frequencies, called “resonance”, is closely associated with the ability of brain regions to oscillate at preferred frequencies ^23,24^. Resonance is generally the outcome of a combination of passive membrane properties of the cell (capacitance and leak conductance) and voltage-gated membrane currents, including the hyperpolarization-activated current (I_h_) ^25^. Since oscillatory activity is a brain-wide phenomenon, it is not surprising that a number of different cell-types show the ability to resonate at subthreshold membrane potentials, albeit at specific preferred frequencies ^23^. In rat hippocampus, pyramidal neurons in *CA1* (r*CA1*) show resonance frequency at 3 – 5 Hz ^25–27^, matching theta wave frequencies in rat (4 – 10 Hz) ^2,20^. In these r*CA1* neurons, it was suggested that resonance could optimize input/output computations from synaptic pathways impinging on the apical dendrites in *CA1*, where I_h_ is particularly prominent ^28^. In addition, r*CA1* pyramidal neurons show a large number of thin, elongated oblique dendrites ^28–30^ and these oblique dendrites possess specialized computational properties ^31–34^. In view of the tight interplay between theta rhythm, resonance frequency and cellular structure/function, it is crucial to determine how concepts and theories based on rodents translate to human hippocampus.

The subcellular structure, biophysical properties and computational capacity of *CA1* pyramidal neurons are well-documented for rodents ^20,35,36^, but these are unknown for human hippocampus. To address this and to test whether reduced theta frequencies in human hippocampus can be explained by cellular function of *CA1* pyramidal neurons, we recorded from *CA1* pyramidal neurons in acute living brain slices of non-pathological human hippocampus obtained during resection surgery. Using detailed compartmental models of h*CA1* pyramidal neurons based on digital reconstructions and physiological recordings, we identify three distinct hippocampal pyramidal neuron types and uncover the consequences of previously unknown morpho-electrical dendritic properties on information routing and nonlinear processing in these large and complex pyramidal neurons that ultimately define human hippocampal function.

## Results

### Human *CA1* pyramidal neurons have large dendritic trees with numerous oblique dendrites

Tissue samples were evaluated for structural abnormalities and only non-sclerotic samples without evident structural alterations (and not being part of deep brain pathology) were included in this study (n = 6 cases). We used small tissue blocks (± 1-2 cm^3^) from human hippocampal bodies for cytoarchitectural assessment, electrophysiological recordings and post-hoc histology (Fig.1). These tissue blocks were obtained during en-block resection of mesio-temporal structures for surgical treatment of drug-resistant epilepsy and in one case to remove a deep brain tumor. NeuN-DAB (see Methods) staining revealed numerous cell-bodies throughout the *Stratum Pyramidale* (*SP*) across canonical regions of hippocampus (Fig.1B). The homogeneous distribution of neuronal somata in human *CA1* (h*CA1*) of the tissue samples confirmed that the tissue was non-sclerotic. All resected samples were evaluated by an expert pathologist and only samples without evident structural alterations were included (Table 1). The thickness of the human *SP* was 1111 μm, 993 – 1232 μm (median, 1^st^-3^rd^ Quartile; n = 6 surgical cases), which not only exceeds the mouse *SP* thickness by a factor of 15 (mouse *SP* 75 μm, ^37^), but also exceeds *SP* thickness of rat (5 cell bodies thick) and macaque (10-15 cell bodies thick) ^31^.

**Fig. 1.**
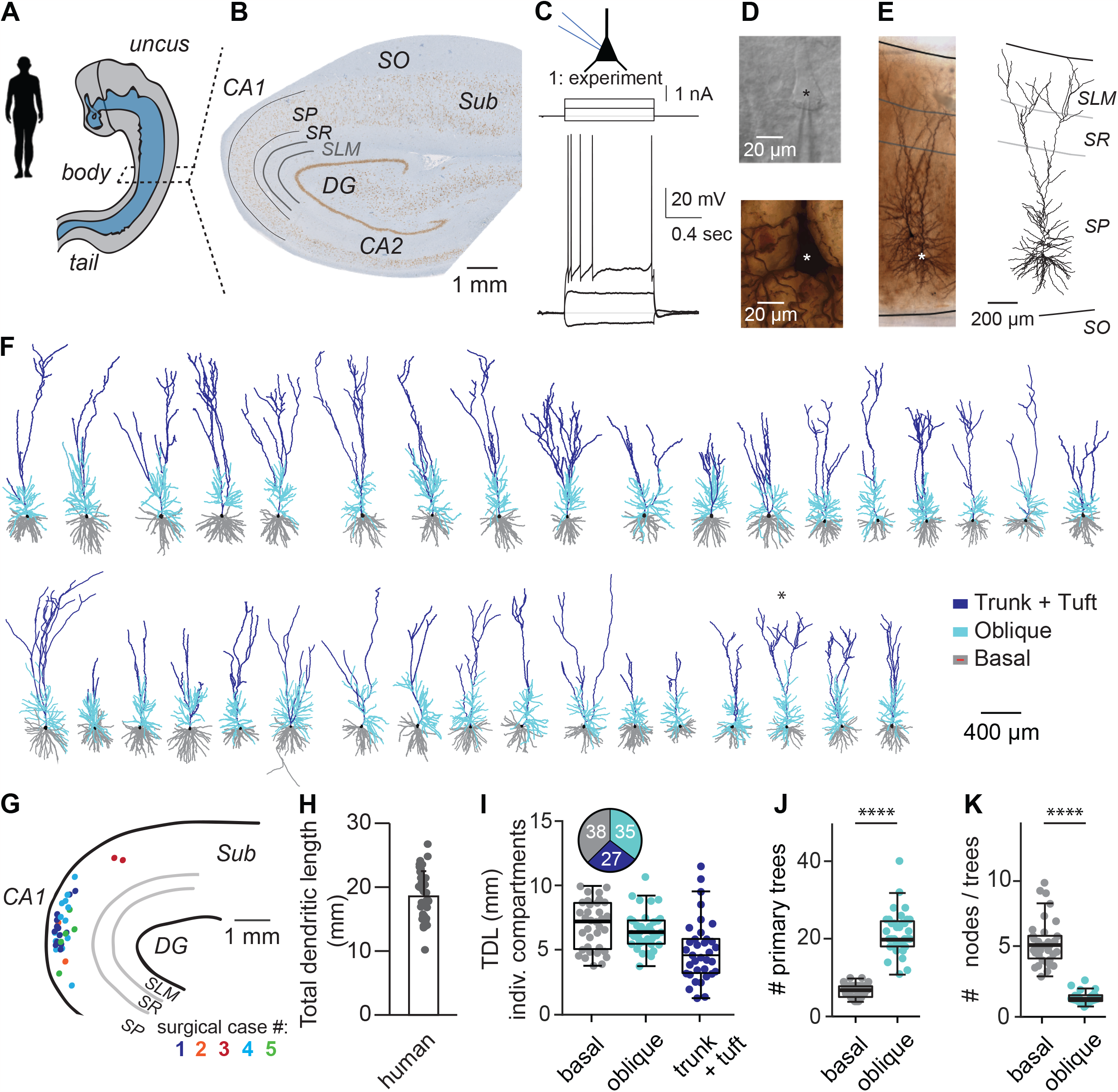
Human CA1 pyramidal neurons have extensive dendritic trees with many oblique dendrites. (A) Human hippocampal tissue originated from comparable parts of the hippocampal body. (B) Cross-section of hippocampus after NeuN histology (see Methods). SO: Stratum Oriens, SR: Stratum Radiatum, SLM: Stratum Lacunosum Moleculare, Sub: Subiculum, CA1, CA2: Cornu Amonis subfield 1 and 2. (C) Experimental current step protocol to quantify passive and active membrane properties. (D, upper panel) Differential interference contrast (DIC) image of an example pyramidal cell body in SP during patch clamp electrophysiology (same cell as in panel C). (D, lower panel) Post-hoc biocytin-DAB stained cell body (same cell). (E, left) Analogous to D (lower panel), but complete pyramidal neuron is illustrated using a collapsed Z-stack image, with layer borders superimposed. Two pyramidal neurons are visible; the example from C – D is indicated by the asterisk. (E, right) Digital reconstruction of the example cell in C – E. (F) Gallery of dendritic reconstructions of hhuman CA1 pyramidal neurons with basal dendrites in gray, oblique dendrites in cyan and main apical trunk and tuft in blue, respectively, soma in black. (G) Annotation of reconstructed morphologies relative to anatomical landmarks and layer position. Colors refer to surgical cases. (H) Total dendritic length (TDL) of human CA1 pyramidal neurons. (I) Compartment specific TDL. Pie chart inset represents the fraction (in %) of individual compartments relative to TDL of full morphology. (J) Number of primary trees is significantly different for basal versus oblique dendrites (basal: 7.0 primary trees, 5.5 – 8.0, obliques: 20.0 trees, 18.5 – 24.5, median, 1^st^-3^rd^ Quartile, n = 35, Mann-Whitney, p < 0.0001). (K) Number of nodes per individual tree is significantly different for basal versus oblique dendrites (basal: 5.0 nodes/tree, 4.1 – 5.7, obliques: 0.9 nodes/tree, 0.7 – 1.2, n = 35, Mann-Whitney, p < 0.0001).

**Table 1:** Surgical cases.

| Case # | Gender | Age (yrs) | Side | Diagnosis | Assessment hippocampal anatomy |
| --- | --- | --- | --- | --- | --- |
| 1 | M | 32 | R | Epilepsy | w/o evident structural alteration |
| 2 | M | 62 |  | No information | No information |
| 3 | F | 23 | R | Epilepsy with tumor | w/o evident structural alteration |
| 4 | M | 49 | R | Epilepsy | w/o evident structural alteration |
| 5 | M | 42 | L | Epilepsy | w/o evident structural alteration |
| 6 | F | 32 | L | Epilepsy | w/o evident structural alteration |

Tissue originated from the hippocampal body, through visual inspection by the neurosurgeon upon resection and anatomical origin was comparable among subjects. We made stable whole-cell patch-clamp recordings of pyramidal neurons (n = 41 recordings, recording duration 30 – 60 minutes, Fig.1C) and probed passive and active membrane properties using a variety of stimulus protocols. Differential interference contrast microscopy indicated the presence of healthy soma’s, based on their appearance (Fig.1D, top) with an average diameter of 19 ± 3 μm and average surface area of 330 ± 60 μm^2^ (n=32), matching previously reported soma size from healthy subjects ^37^. Recorded neurons were dye-filled with biocytin for post-hoc histology (Fig.1D, bottom), and digital reconstruction (Fig.1E right). Additionally, cytoarchitectural layers were determined to annotate reconstructed neurons to anatomical landmarks (Fig.1E right, Suppl.Fig.1, see Methods). Thus, hippocampal slices contained healthy neurons with stable, intact membranes and neurons showed repetitive action potential spiking in response to current injection. We did not observe spontaneous epileptiform activity in slices.

The tissue samples used for patch-clamp electrophysiology contained intact *CA1, CA2, CA3* regions in addition to *Dentate Gyrus (DG)* and at least part of *Subiculum* (Fig.1B). Within the region of interest of the current study (i.e. *CA1*), samples always included all canonical, cytoarchitectural layers of hippocampus *CA1* (Fig.1B, 1E, Suppl.Fig.1). Anatomical landmarks (*DG*) and layer borders subsequently allowed the annotation of n = 35 individual reconstructed neurons to a standardized framework, including cellular position within the *Stratum Pyramidale (SP)* with respect to the *DG* apex and radial position in *SP* relative to *Stratum Oriens* (*SO*) and *Stratum Lacunosum Moleculare* (*SLM*) borders (Fig.1F, G, Suppl.Fig.1).

The reconstructed h*CA1* pyramidal neurons showed extensive dendritic trees that were relatively intact (Fig.1F). The total dendritic length (TDL) of single h*CA1* pyramidal neurons accumulated to 18.6 ± 3.9 mm (average ± stdev, n = 35, Fig.1H). This significantly exceeds TDL in rat (12.9 ± 1.5 mm, n = 29, ^38,39^, hamster (3.0 ± 0.6 mm, n = 66, ^40^ or mouse (3.7 ± 0.6 mm, n = 46^37^, ANOVA, p < 0.001). Thus, human *CA1* pyramidal neurons have extended dendrites compared to rodent species.

Next, we quantified the length of basal, oblique and apical trunk + tuft dendrites of h*CA1* pyramidal neurons separately as these subtrees have been reported to contribute to specific biophysical and computational properties in rodent *CA1* pyramidal neurons ^33,34,41–43^. We find that these 3 major dendritic compartments contribute equally to the total dendritic architecture (Fig.1I, inset).

Oblique dendrites may be of particular importance to computational properties of individual neurons ^33,34,44^. We thus quantified the structural properties of basal vs oblique dendrites in h*CA1* pyramidal neurons. We found that the number of primary trees is 3-fold larger for basal dendrites compared to oblique dendrites (p < 0.0001, Mann-Whitney test, Fig.1J). Oblique dendrites showed much less branching per tree compared to branching in basal dendrites (p < 0.0001, Mann-Whitney test, Fig.1K).

### I_h_ currents drive resonance at human theta oscillation frequencies

Given that no previous reports on whole-cell biophysical properties exist for non-pathological human *CA1* pyramidal neurons, we assessed basic neurophysiological properties of the neurons (Fig.2A). First, intrinsic resting membrane potentials were -63.4 mV (-66.2 to -60.1 mV, median, 1^st^ Quartile - 3^rd^ Quartile range) and we observed spontaneous action potential (AP) firing in only 3 out of 41 recordings (from n = 3 different slices and 2 surgical cases) and only immediately after establishing whole-cell configuration. No spontaneous APs were observed during subsequent protocols confirming that the epileptic focus was not within our hippocampal slices. In addition, the input resistance was 37.8 MΩ (33.7 – 58.9 MΩ, n = 41) (median, 1^st^ – 3^rd^ Quartile, Fig.2B) and the membrane time constant (tau) was 21.7 ms (16.2 – 29.6 ms, n = 41) (median, 1^st^ – 3^rd^ Quartile).

**Fig. 2.**
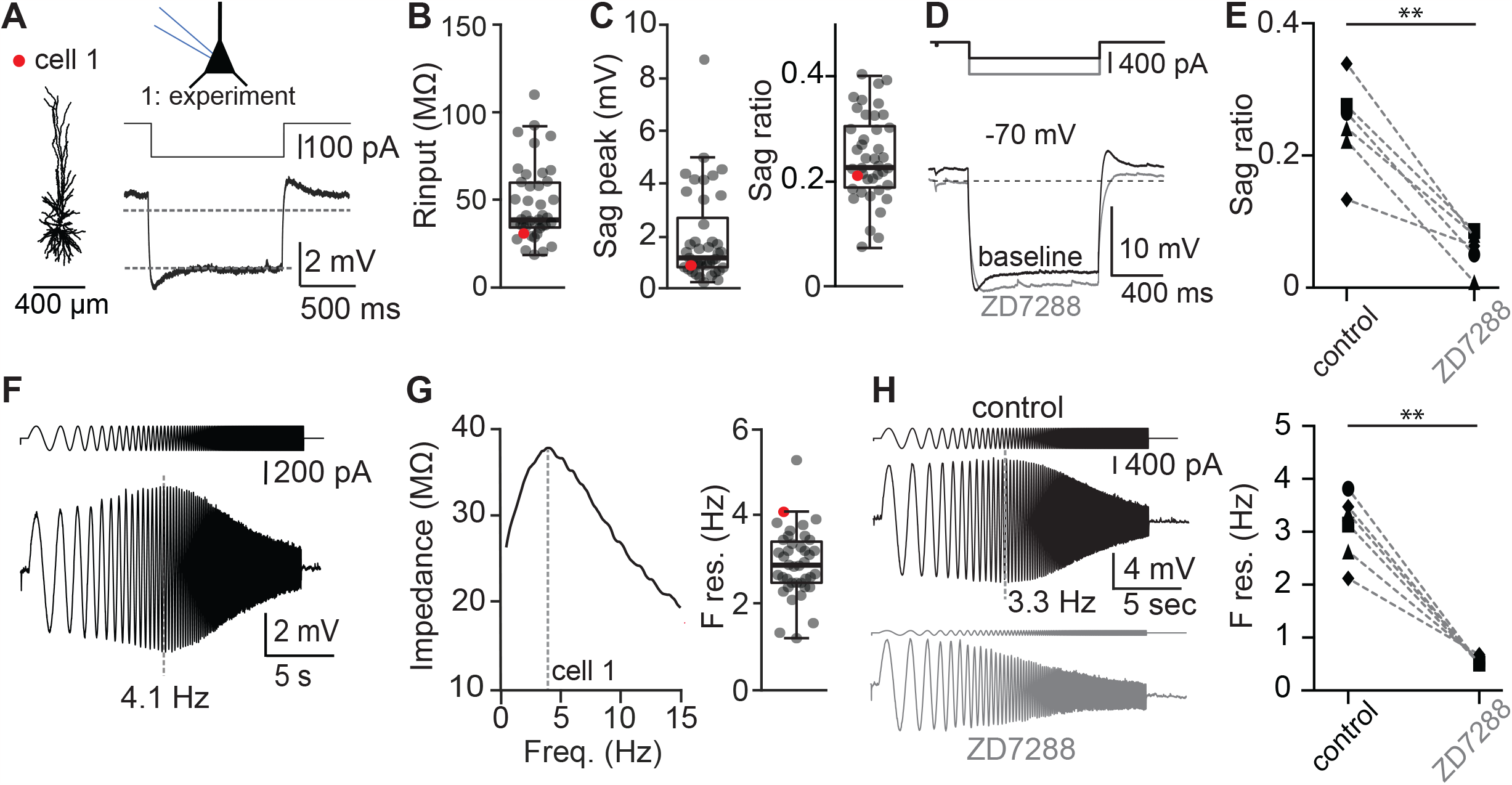
I_h_ currents drive resonance at human theta oscillation frequencies. Example hCA1 pyramidal neuron with digital reconstruction of dendritic morphology and hyperpolarization current to uncover I_h_ (or sag current). Dashed lines correspond to -70 mV and steady-state hyperpolarization (close to -73mV, see Methods), respectively. (B) Population statistics for input resistance. (C, left) Population statistics for sag peak and (C, right) sag ratio. (D) ZD7288 (HCN channel antagonist) blocks sag current during hyperpolarizing current injections (dark gray: control, light gray: ZD7288). (E) ZD7288 significantly reduces sag ratio across regions, (n = 6, Wilcoxon signed-rank, p < 0.01). Symbols refer to entorhinal cortex (EC, •), temporal cortex (T, ◼), hippocampus (H, ♦) or frontal cortex (F, ▾). (F) Chirp protocol to extract resonance properties (see Methods). (G, left) Impedance profiles of example cell #1 in response to the chirp input current. (G, right) Population statistics for preferred frequency (ie. resonance frequency, n = 40, 2.9 Hz, 2.5 - 3.4 median, 1^st^ – 3^rd^ Quartile). (H, left) ZD7288 abolishes resonance properties in response to chirp protocol (black: control, gray: ZD7288). (H, right) ZD7288 significantly blocks resonance properties across regions (n = 6, Wilcoxon signed-rank, p < 0.01). Symbols same as (E).

Rodent *CA1* pyramidal neurons show hyperpolarization-actived I_h_ currents, which endow them with resonance properties ^25,26,45–47^. This has never been tested in h*CA1* pyramidal neurons. We measured the voltage response to hyperpolarizing current injection to quantify the properties of I_h_ (Fig.2A), indicative of HCN channel activity. The amplitude of the hyperpolarizing current was scaled to the input resistance to consistently generate a hyperpolarization to approximately -73 mV. A sag response was found in all recordings (1.2 mV, 0.8 – 2.4 mV, median, 1^st^ – 3^rd^ Quartile, n = 41, Fig.2C, left). Normalization to the steady-state voltage response generates the dimensionless sag ratio, which was 0.23, 0.19 – 0.30 (median, 1^st^ – 3^rd^ Quartile, Fig.2C, right). We confirmed that this sag is due to HCN channel activation as it was consistently blocked by the HCN channel antagonist ZD7288 (Fig.2D, E).

To test whether h*CA1* pyramidal neurons show resonance and if so, at what frequency, we measured the voltage response to a chirp stimulus (Fig.2F, see Methods, for mouse data see Suppl.Fig.2). When voltage response amplitudes showed a maximum at frequencies above 1Hz, neurons were considered to be resonant (Fig.2G). Across recordings, we found that 40 out of 41 recorded neurons showed resonance frequencies above 1 Hz (98%). On the population level, the preferred frequency of h*CA1* pyramidal neurons was 2.9 Hz, 2.5 – 3.4 Hz (n = 40, median, 1^st^ - 3^rd^ Quartile, Fig.2G, right) with consistent values across surgical cases (F res. case #1 - #5, F res. 2.5 Hz (n=13), 2.8 Hz (n=3), 3.9 Hz (n=1), 3.2 Hz (n=15), 2.8 Hz (n=8). Resonant properties were consistently abolished by the HCN channel antagonist ZD7288 across different cortical regions (Fig.2H).

### Human *CA1* contain three morpho-electric pyramidal neuron types

To determine whether separate groups of human *CA1* pyramidal neurons with distinct morphological or biophysical properties could be identified, we analyzed a large morpho-electric parameter space of recorded pyramidal neurons. We used anatomical location along the dorsal-ventral axis of the *CA1 Stratum Pyramidale* (*SP*), normalized as a fraction relative to the SP-SLM dimension to account for variability in *SP* thickness between surgical cases. Both passive properties (see Fig.2) as well as five active properties of action potentials (APs) were used, which were extracted from the first AP at rheobase (see Methods, Fig.3A). Morphological measures included (amongst others) total dendritic length, # of obliques, # of nodes and # of bifurcations in the first 200 micrometer of the apical tree (i.e., “early bifurcations”, Fig.3B ^37^.

**Fig. 3:**
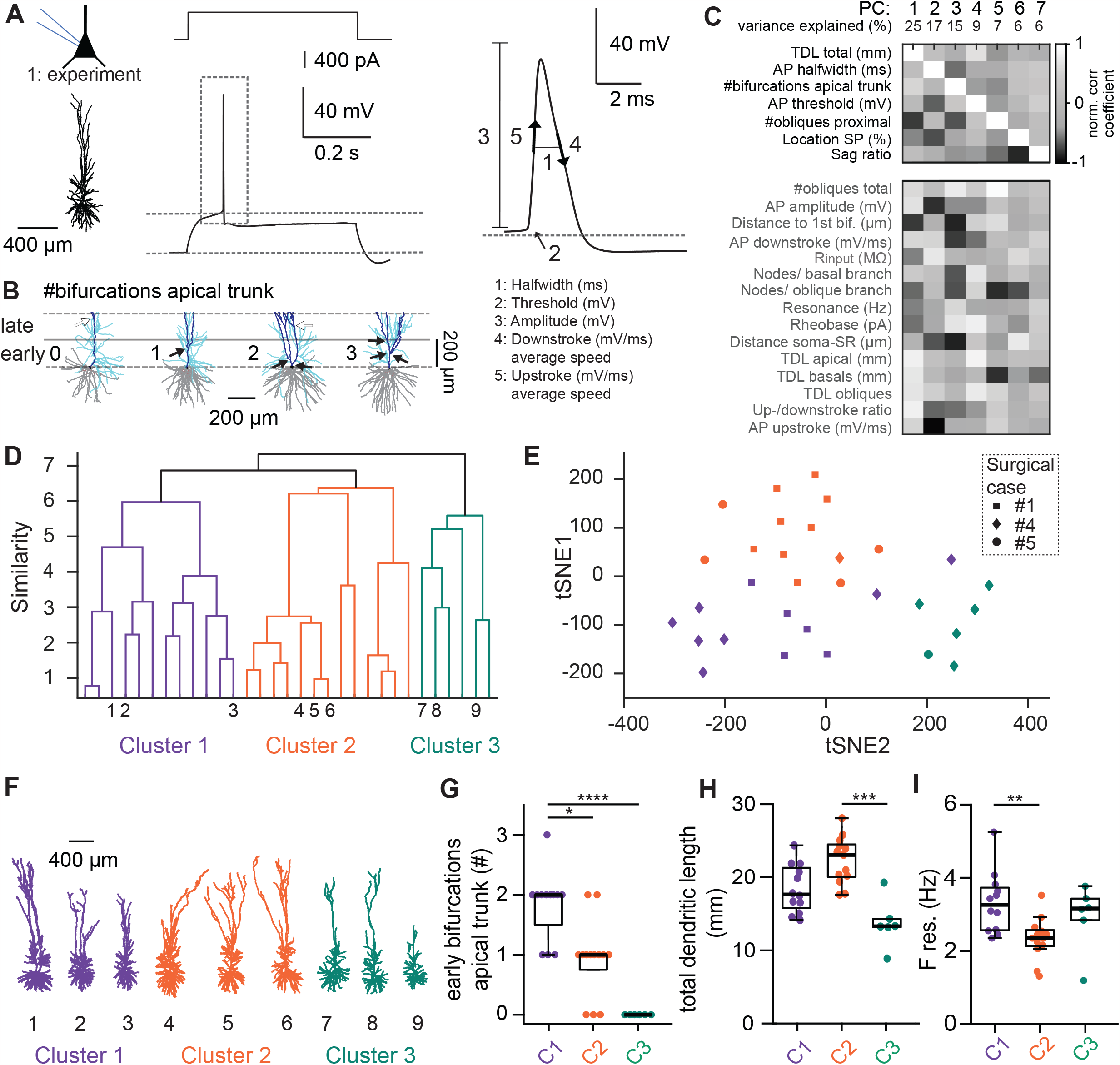
Human CA1 contains three types of pyramidal neurons with distinct morpho-electric properties. (A, left) Example morphology with matching positive current injection at rheobase. (A, right) Action potential (AP) features quantified. (B) Four example reconstructions showing ‘early’ apical trunk branching within the first 200 micrometer of the main apical trunk (zero, one, two or three bifurcations, respectively (see also (38)). Note that neurons can also contain ‘late’ bifurcations, located > 200 micrometers from apical origin. (C) The morphological and electrophysiological features included for PCA. Gray values indicate the normalized correlation coefficient for each individual feature versus the principal component (see Methods). Features dominating the principal components are listed on top, ranked according to PC weight (white square). Additional features (listed below) are listed without weight. (D) Unsupervised hierarchical cluster analysis based on features in (C). Small numbers adjacent to the horizontal axis refer to example morphologies. (E) t-SNE plot for the three main cell classes (symbols refer to surgical case). (F) Example morphologies for the 3 main cell classes. (G) The number of early bifurcations is significantly different between main cell classes. Note: cluster 3 neurons completely lack early bifurcations in the apical trunk. (H) Total dendritic length is highest in cluster 2 neurons. (I) Resonance frequency is lowest in cluster 2 neurons. Statistics: Kruskal-Wallis (unpaired, non-parametric ANOVA) with Dunn’s post-hoc test, * p < 0.05, ** p < 0.01, *** p < 0.001).

We used principal component analysis on 22 morpho-electric features to determine which principal components (PCs) dominated and how each feature contributed to the PCs (Fig.3C). In short, only principal components that explained more than 5% of the variance were included in the dataset, resulting in a total of 7 principal components, which together explained approximately 85% of the variance. We subsequently determined which parameters contributed most to the 7 PC’s, reflected as correlation coefficient normalized to the feature with maximal contribution (Fig.3C).

Unsupervised hierarchical cluster analysis distinguished three main classes of h*CA1* pyramidal neurons (Fig.3D, see Methods). The t-distributed stochastic neighbor embedding (t-SNE) plot showed groupings of neurons with shared morpho-electric properties that clustered together (Fig.3E). Each cluster contained neurons from multiple surgical cases. The most obvious differences between these 3 h*CA1* cell classes were i) the number of early bifurcations in the apical trunk, ii) dendritic architecture, measured as total dendritic length and iii) resonance properties (Fig. 3G - 3I). In short, Cluster 1 neurons show highest number of early bifurcations in the apical trunk and intermediate values for total dendritic length and resonance frequency. Cluster 2 neurons have largest total dendritic length and lowest resonance frequency. Cluster 3 neurons show no early apical bifurcations, lowest total dendritic length, and intermediate resonance frequency.

We also found a difference in action potential waveform (Suppl.Fig.3), but no differences in total number of oblique dendrites, action potential amplitude or sag ratio. *SP* location did significantly vary between main classes (Suppl.Fig.3), but distributions showed considerable overlap between populations. Thus, human *CA1* contains at least three distinct morpho-electric types of pyramidal neurons with distinct properties that intermingle in human SP.

### Oblique dendrites act as independent processing compartments

To investigate the impact of structural and electrophysiological properties of human *CA1* pyramidal neurons on the processing of synaptic events on their computational properties, we built biophysically detailed compartmental models of these neurons. We constructed experimentally-based compartmental models for n = 16 individual h*CA1* neurons, sampled from each of the three cell classes described above (Fig.3, Fig.4 and Methods). Results of fitting cable parameters to these cells are exemplified in Fig.4B-C. Complete cable parameters per cell can be found in Table S3. We performed an extensive parameter search for optimal fitting the model to the experimental data, for all 16 modeled neurons. The models were fitted to both responses to hyperpolarizing current steps (to model HCN channel activation), and to responses to chirp protocols (Fig.4C). The resulting preferred frequency accurately reproduced the experimental data (experiment: 2.35 Hz, simulation: 2.37 Hz, Fig.4C, Table S3). The similarity between experimental data and simulation is highly non-trivial emphasizing that the optimization procedure used accurately incorporated crucial biophysical parameters that closely reflect experimental data.

**Fig. 4.**
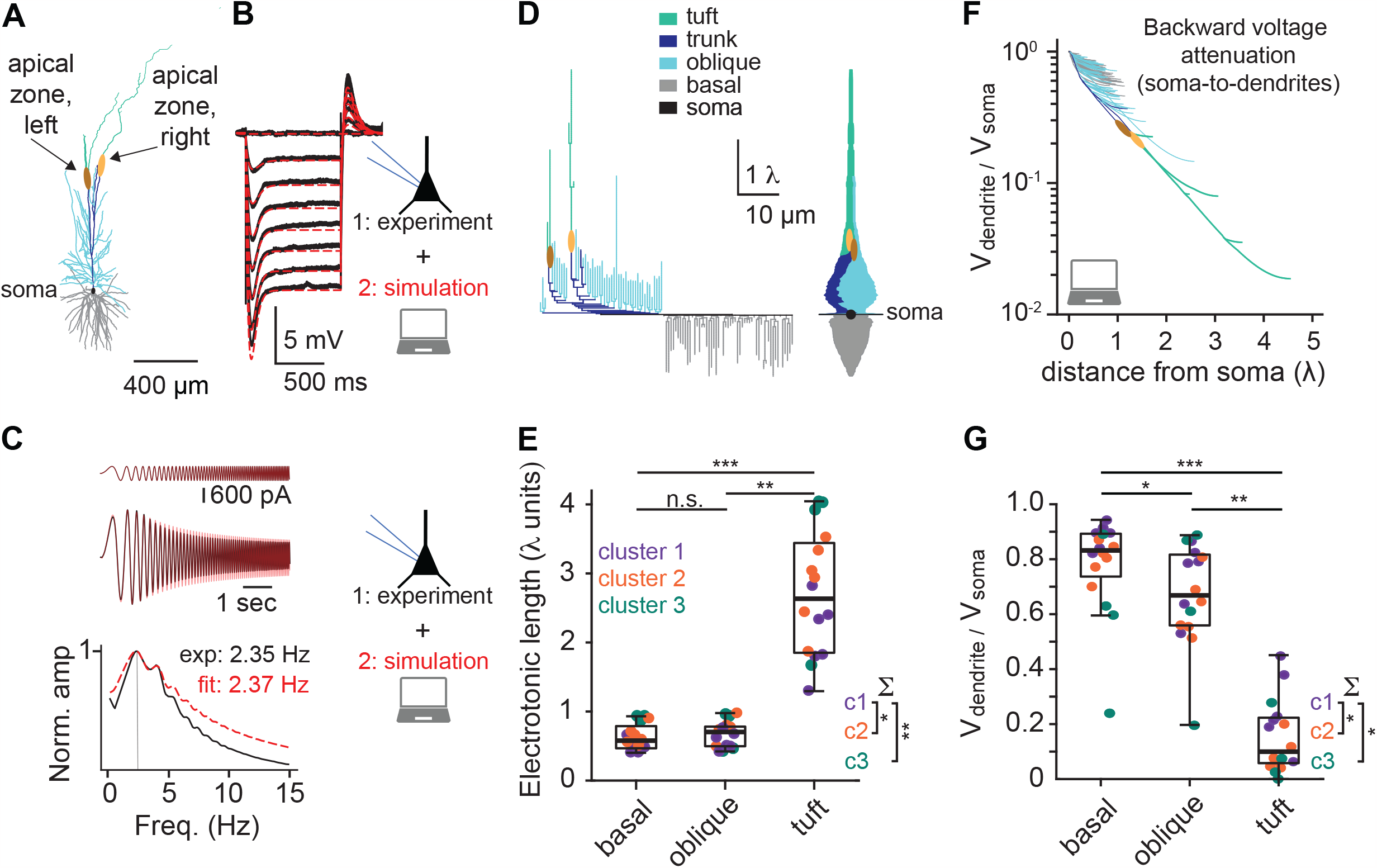
Cable properties of human CA1 pyramidal neurons. (A) Example morphology used for biophysical modeling (identity: cluster 2), including two apical zones to read-out the propagation of electrical currents in the apical dendrite. (B) Experimental data (black) and model fit (red) of hyperpolarizing current steps applied to the example morphology shown in (A, -0.7 to 0.0 in 0.1 nA steps). The extracted passive parameters for this example morphology are: R_m_ = 149,994 Ωcm^2^, C_m_ = 0.7 μF/cm^2^, R_a_ = 300 Ωcm. This model included spatially nonuniform I_h_ conductance (see details in Methods). (C) Experimental data (black) and model fit (red) of resonance properties. (D) Left. Dendrogram in cable units for the modeled pyramidal neuron shown in (A), based on the extracted cable parameters. Right. “Equivalent cable” of neuron in panel A. Soma is depicted by the black circle, the relative contribution of each part of the dendritic tree is illustrated by the corresponding colors. (E) Population statistics for electrotonic length (in cable units) for n = 16 models. (F) Simulation of steady state voltage attenuation from the soma to the dendrites. (G) Population statistics for steady state voltage attenuation (n = 16). Note: residual signal in the tuft is only 10% of the original amplitude at the level of the soma. Statistics between dendritic compartments: Friedman (paired, non-parametric ANOVA) with Dunn’s post-hoc test, * p < 0.05, ** p < 0.01, *** p < 0.001. Statistics for class-specific properties: Kruskal-Wallis (unpaired, non-parametric ANOVA) with Dunn’s post-hoc test, * p < 0.05, ** p < 0.01.

From the soma viewpoint, the basal and oblique dendrites in the example neuron were electrotonically compact (low λ values), whereas the distal apical tuft was electrically decoupled from the soma (high λ values, Fig. 4D). The particularly large membrane area of the oblique dendrites in h*CA1* pyramidal neurons is reflected by the large increase in the diameter of the equivalent cable for these neurons (Fig.4D right, cyan, see Methods). This implies that the oblique dendrites impose a large conductance load (“current sink” resulting in electrical decoupling) for signal transfer between the soma and the apical tuft (and vice versa). For the population (n = 16 models), we find that the electrotonic length from soma to dendritic tips (in cable units, λ) is comparable for oblique and basal dendrites and largest for the apical tuft (Fig.4E, Friedman paired test, Dunn’s post-hoc p < 0.01 or p < 0.001). This resulted in a steep voltage attenuation from the soma to the apical tuft, even in steady state conditions (Fig.4F). The backwards voltage attenuation (soma to dendrites) was highly specific for the different dendritic compartments; basal, oblique and tuft dendrites (Fig.4G, median basal: 0.83, obliques: 0.67, tuft: 0.10, Friedman paired test, Dunn’s post-hoc p < 0.05, p < 0.01 or p < 0.001).

When we compared electrotonic length and attenuation for the three main cell classes of human *CA1* pyramidal neurons, we found class-specific properties. After generalization for dendritic tree compartment, electrotonic length was smaller for cluster 1 neurons compared to cluster 2 and cluster 3 neurons (p < 0.05, Dunn’s post-hoc test). Soma-to-dendrites voltage attenuation was less pronounced in cluster 1 neurons compared to cluster 2 and cluster 3 neurons (p < 0.05, Dunn’s post-hoc test, Fig.3 inset).

Due to steep antidromic voltage-attenuation of somatic electric activity, we hypothesized that the oblique and distal apical dendrites can process subthreshold synaptic information relatively independent from the soma. To test this, we first studied signal transfer from different dendritic compartments to the soma and other dendritic compartments. The reconstructed neuron in Fig.5A shows the distinct dendritic compartments in h*CA1* pyramidal neurons: basal (gray), proximal (blue) and left and right distal oblique dendrites (red and green, respectively). We simulated sinusoidal input currents across different frequencies (0.01 – 1000 Hz) at different oblique dendrite tips and measured the membrane voltage response at three locations indicated in the reconstructed neuron in Fig.5A: 1) the left distal apical zone (Fig.5B), 2) the right distal apical zone (Fig.5C) and 3) the soma (Fig.5D). The injected currents at all dendritic locations had largest impact on subthreshold membrane potential changes at frequencies around 3 Hz (Fig.5B-D). However, there was a large difference in voltage response amplitudes, depending on the dendritic location of current injections. At the left apical zone on the distal dendrite, current injections in the left oblique dendrites (red) had strongest impact on the membrane potential changes there, whereas injections in other dendritic compartments (right obliques in green, proximal dendrites in blue, basal dendrites in gray) had much less impact on membrane potential changes in the left apical zone (Fig.5B). This pattern was mirrored for the right apical zone (Fig.5C), with distal right oblique dendrites being most effective. At the soma, current injections in basal and proximal dendrites had the strongest impact on membrane potential changes, whereas current injections in the distal obliques had only modest impact on membrane potential changes (Fig.5D).

**Fig. 5.**
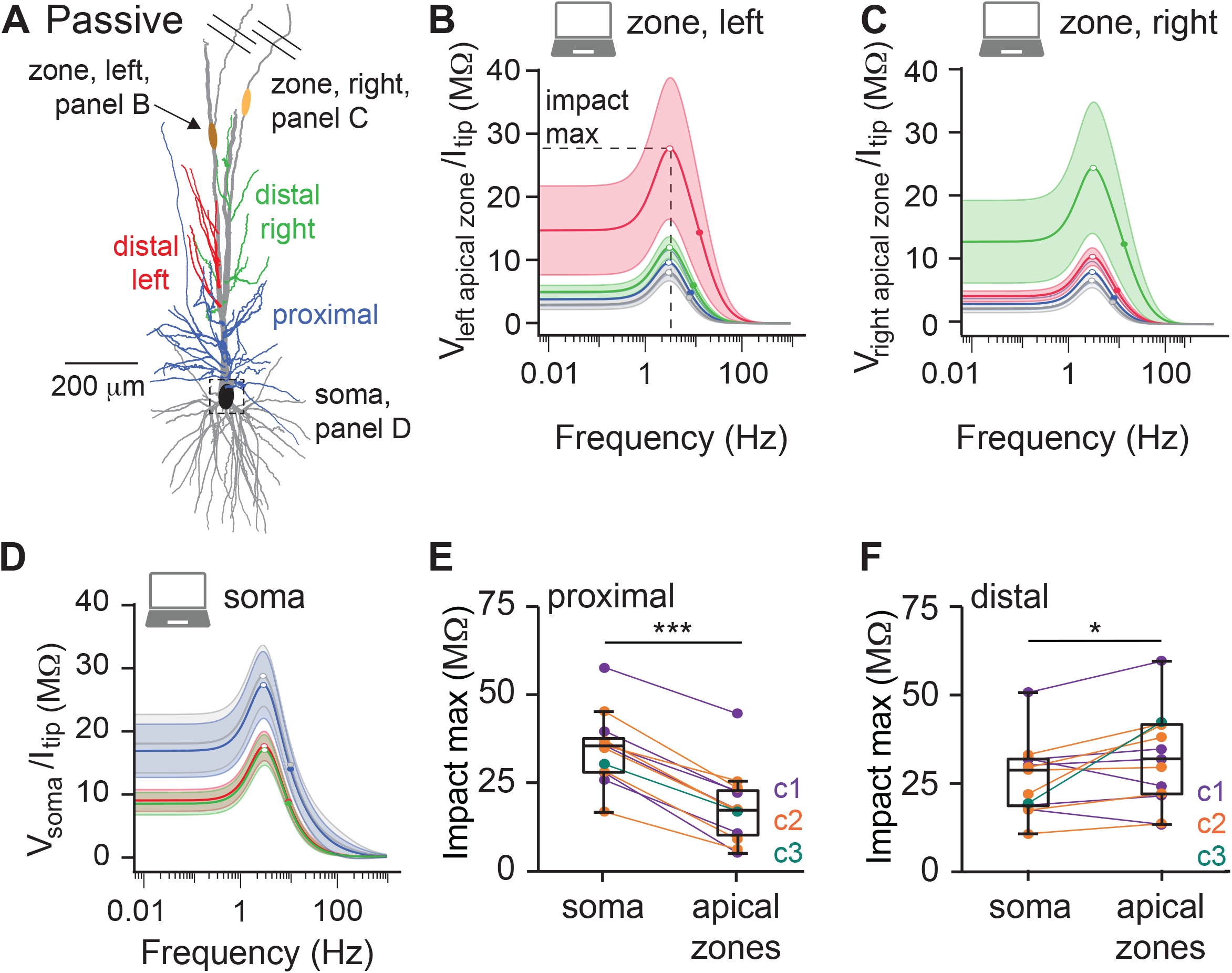
Preferential routing of subthreshold inputs in proximal and distal dendrites. (A) Example morphology with oblique dendrites color coded as a function of location relative to the early apical trunk bifurcation, including two apical zones to read-out the propagation of electrical currents in the apical dendrite. Oblique dendrites proximal to the bifurcation in blue, oblique dendrites distal to the bifurcation in red and green for left and right apical branches, respectively. (B) Frequency dependent impact on the left apical zone in response to sinusoidal currents generated in different dendritic compartments. Effect of current injected at distal, left oblique dendrites is shown in red, effect of distal right obliques in green, proximal obliques in blue and basal dendrites in gray. (C) Analogous to (B) but the effect of injected current on the right apical zone. Note that current injection in distal, but left oblique dendrites (red curve) has reduced impact relative to current injection in distal, right oblique dendrites (green curve). (D) Analogous to (B, C) but the effect of injected current on the soma. (E) Population statistics for ‘impact max’ shown in B-D indicating that current injection in proximal dendrites has more impact on membrane potential in the soma compared to apical zones. Color code of individual data points match cell cluster identity (see Fig.3). (F) Analogous to (E) but the impact of current injection in distal oblique dendrites on soma or apical zones. Note that distal oblique dendrites have more impact on apical zones relative to soma. Statistics: Wilcoxon (paired, non-parametric), * p < 0.05, *** p < 0.001.

Signal transfer from the left and right oblique dendrites to the apical zone on their own side was more effective compared to signal transfer to the opposite side (Suppl.Fig.4, Friedman with Dunn’s post-hoc test, p < 0.001). Most importantly, signal transfer from the proximal oblique dendrites to the soma was significantly more efficient compared to signal transfer from proximal oblique dendrites to the apical zones (Fig.5E, Wilcoxon paired test, p < 0.001). In contrast, signal transfer from the distal oblique dendrites was significantly more efficient towards the apical zones compared to signal transfer from the distal oblique dendrites to the soma (Fig.5F, Wilcoxon paired test, p < 0.05). We did not find differences between main cell classes in signal transfer for proximal or distal oblique dendrites to soma or apical zones. In summary, passive signals are processed highly asymmetrically in the large dendritic trees of human *CA1* pyramidal neurons and translates to preferential routing towards soma or into the apical dendritic tree, depending on their location of origin.

### Active signal propagation in dendrites distributes non-linearly

Both rodent and human neocortical pyramidal neuron dendrites show active electrogenesis, generating sodium and calcium-based action potentials ^48–54^. Simultaneous recordings from soma and dendrite have shown that electrical activity recorded in the dendrite is causally linked to changes in the amplitude of the after-depolarization (ADP) of the action potential, recorded at the soma ^48^. Blocking voltage-gated calcium channels (VGCC) prevents dendritic electrogenesis as well as the somatic ADP, both in rodent and human neocortical pyramidal neurons ^48,49^. This could suggest that also in human pyramidal neurons, the ADP reflects dendritic electrogenesis. Human *CA1* pyramidal neurons also showed a pronounced ADP (Fig.6A) that increased in amplitude when the frequency of successive action potentials increased (Fig.6B). To validate whether the ADP was carried by VGCC activity, we washed in the VGCC blocker cadmium (CdCl_2_, Fig.6C). This reduced the amplitude of the ADP at all successive AP frequencies tested (Fig.6D), suggesting that hCA1 pyramidal neurons exhibit VGCC-dependent electrogenesis ^55^.

**Fig. 6:**
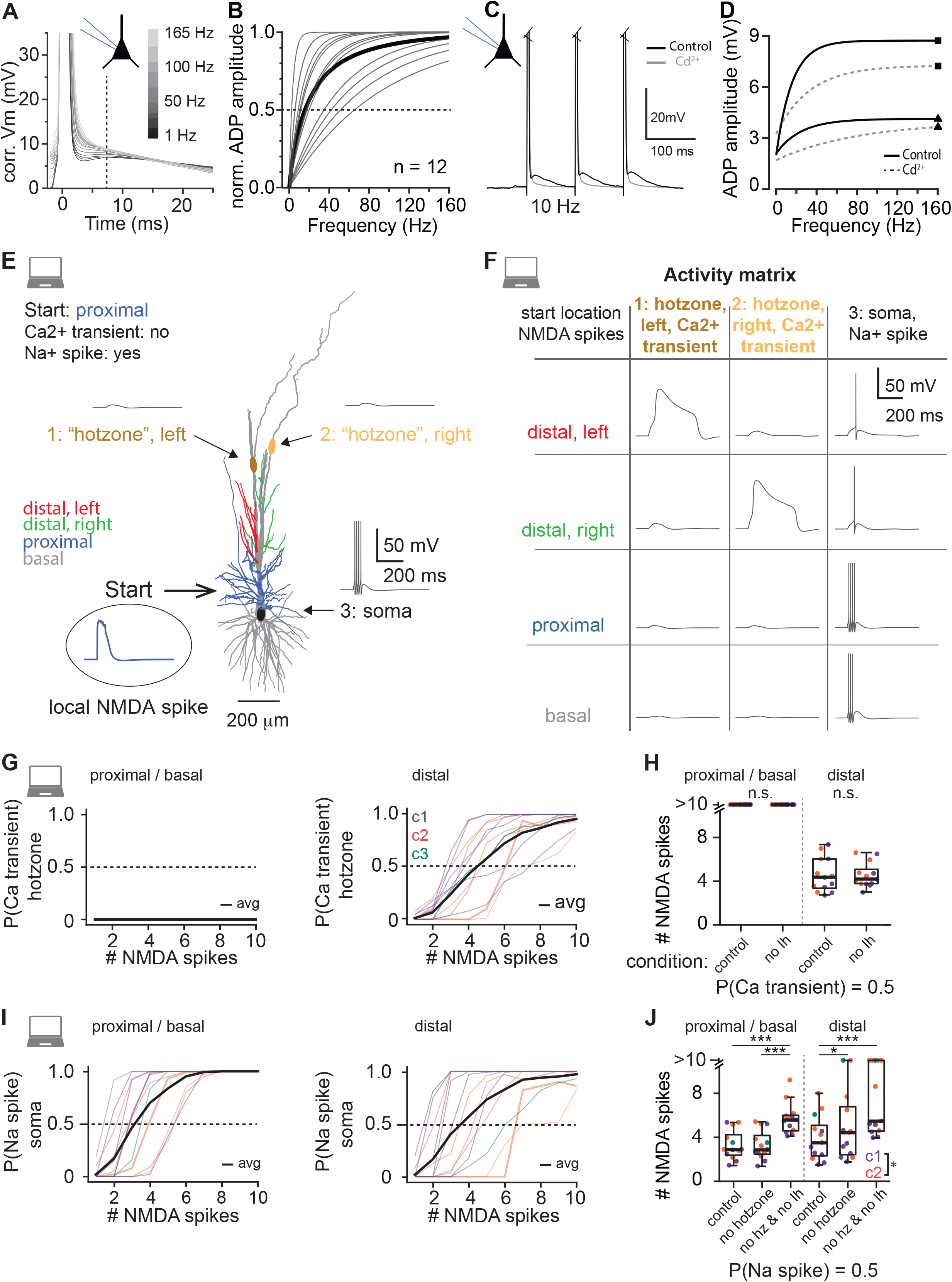
Dendritic I_h_, local NMDA spikes and Ca^2+^ hotzones shape non-linear output spiking. (A) Experimental voltage traces of an example neuron showing the action potential (AP) after-depolarization (ADP) recorded at the soma. Three APs were elicited at a controlled frequency, but only the last APs are visualized. (B) Normalized ADP amplitude as a function of AP triplet frequency. Individual gray lines represent exponential fits for individual recordings (n = 12), the population average is shown in black (bold). (C) Wash-in of Cd^2+^ reduces the ADP amplitude, indicative of contribution of voltage dependent Ca^2+^ channels to AP waveform. (D) Frequency-dependent increase in ADP amplitude is reduced by Cd^2+^. (E) Example model (same as Fig.4, 5) with apical zones enriched with voltage-dependent calcium channels (now: “hotzones” and the soma enriched with voltage-dependent sodium and potassium channels. Voltage traces of the simulation illustrate the response to 7 NMDA spikes evoked in proximal oblique dendrites. Note that NMDA spikes in proximal obliques trigger somatic Na^+^ spikes, but fail to trigger apical Ca^2+^ transients in the hotzones. (F) Activity matrix to illustrate the efficiency of different types of oblique dendrites to trigger apical Ca^2+^ transients or somatic Na^+^ spikes. (G, left) The probability of activating a Ca^2+^ transient in one of the hotzones as a function of the number of NMDA spikes in proximal oblique or basal dendrites. Note that these dendritic compartments fail to trigger Ca^2+^ transients in the apical hotzones. (G, right) Analogous to (G, left) but for distal oblique dendrites. Thin lines represent individual biophysical models (color coded by cluster identity), thick black line represents population average. (H) Population statistics for the simulations in (G). Removing I_h_ from the biophysical models did not affect hotzone excitability. Statistics: Wilcoxon (paired, non-parametric), n.s.: not significant. (I) Analogous to G, but the probability of activating a somatic Na^+^ spike as function of the number of NMDA spikes in proximal oblique/basal or distal oblique dendrites. (J) Population statistics for the simulations in I (Friedman, p < 0.05, or p < 0.001). Note that removing the hotzones from the biophysical models reduces excitability and thus increased the number of required NMDA spikes to trigger a Ca^2+^ transient in the hotzones, but only for the distal oblique dendrites. Removal of I_h_ also increased the number of required NMDA spikes to trigger a somatic Na^+^ spike, but in a dendrite specific manner. Note that cluster 2 neurons are generally less excitable (P_Na+_ = 0.5 for C1: 2.5, 1.8 – 2.9 NMDA spikes, C2: 4.1, 2.9 – 5.2 NMDA spikes, median, 1^st^ – 3^rd^ Quartile, Mann-Whitney, p < 0.05).

Since we found that subthreshold synaptic inputs are non-linearly processed by the large dendritic trees of the human *CA1* pyramidal neurons, we asked whether dendritic generation and propagation of sodium and calcium-based action potentials also showed non-linear properties. We thus expanded the individual neuron models by including voltage-gated calcium channels in the apical dendritic tree to support active dendritic calcium transients in the apical tree (i.e. apical zones now have active properties, hence “hotzones”, ^48^). In addition, we created an active soma through voltage-gated sodium and potassium channels (see Methods).

Human neocortical pyramidal neurons can generate NMDA spikes ^56^. To simulate excitability under biologically realistic conditions, we generated local NMDA spikes in different dendritic compartments and tested how they induced calcium-based plateau potentials and sodium-based action potentials (calcium transients and sodium spikes, Fig.6E, 6F, Suppl.Fig.5). For instance, NMDA spikes in proximal oblique dendrites triggered a Na^+^ spike in the soma, but Ca^2+^ transients were not generated in either of the apical hotzones (Fig.6E). NMDA spikes in distal oblique dendrites revealed a different input/output function: NMDA spikes triggered a single Na^+^ spike in the soma and a Ca^2+^ transient in the distal hotzone, but only when located on the same side of the stimulated oblique dendrites (Fig.6F).

NMDA spikes in proximal and basal dendrites generally failed to trigger calcium transients in the distal hotzones, even when 10 NMDA spikes were triggered across basal and proximal oblique dendrites (Fig.6G, i.e. [P]robability = 0). In contrast, the probability of activating a Ca^2+^ spike in one of the hotzones increased as a function of the number of NMDA spikes generated simultaneously in multiple distal oblique dendrites (Fig.6G). Approximately four NMDA spikes across various distal oblique dendrites were sufficient to trigger a Ca^2+^ transient in 50% of the trials (Fig.6G, i.e. [P]robability = 0.5). Removing I_h_ from the model did not change the number of required NMDA spikes to trigger a Ca^2+^ transient in the distal dendrites (Fig.6H; Mann-Whitney, p > 0.05), suggesting that this property results from morphological properties and passive cable properties rather than I_h_ channel activity.

NMDA spikes in proximal oblique dendrites were effective at triggering Na^+^ spikes at the level of the soma (Fig.6I, J) as approximately 3 NMDA spikes triggered somatic sodium spikes in 50% of the trials (i.e. P = 0.5). NMDA spikes in distal oblique dendrites were less effective in generating somatic sodium spikes and required ± 4 NMDA spikes to trigger somatic sodium spikes in half the trials (Fig.6I, J). Removing the Ca^2+^ hotzones from the apical dendrite did not affect the number of NMDA spikes required to trigger sodium spikes from the proximal obliques and basal dendrites. However, subsequent removal of I_h_ from the model did increase the number of NMDA spikes required to trigger somatic sodium spikes (Fig.6J). In contrast, removing the Ca^2+^ hotzones from the apical dendrite increased the number of NMDA spikes required on distal oblique dendrites to trigger somatic sodium spikes, and this was further augmented by removing I_h_ from the model (Friedman with Dunn’s post hoc test, p < 0.05, p < 0.001; Fig.6J). Thus, generation and active propagation of calcium transients and sodium spikes in human *CA1* pyramidal neurons depend asymmetrically and nonlinearly on the activation of specific subgroups of dendritic branches. In addition, we find that pyramidal neurons from cluster 2 need a larger number of NMDA spikes to trigger a Na^+^ spike compared to cluster 1 neurons (Mann-Whitney, p < 0.05). This difference persists after removing the Ca^2+^ hotzones (Mann-Whitney, p < 0.05) and after removing both the hotzones and I_h_ (Mann-Whitney, p < 0.001).

In summary, our data-driven modeling shows asymmetric passive and active signal transfer in the large human *CA1* pyramidal neurons (Fig. 7A, B). More specifically, the extensive branching and extended cable of the dendritic tree of these neurons combined with their large number of oblique dendrites implies that many dendritic compartments can act independently from each other. To translate this into computational capacity, we used a two-layered model for the three main classes of human *CA1* pyramidal neurons separately, considering differences in dendritic architecture and total dendritic length (Fig.7C). We find that computational capacity is highest for cluster 2 neurons, even though cluster 1 neurons have most dendritic subunits (i.e. individual dendritic trees). Cluster 3 neurons have lowest computational capacity, reflecting their relatively simple dendritic structure (Fig.3). Compared to mouse, computational capacity of human *CA1* pyramidal neurons is 8-fold (cluster 3) to 12-fold (cluster 2) larger.

**Fig. 7:**
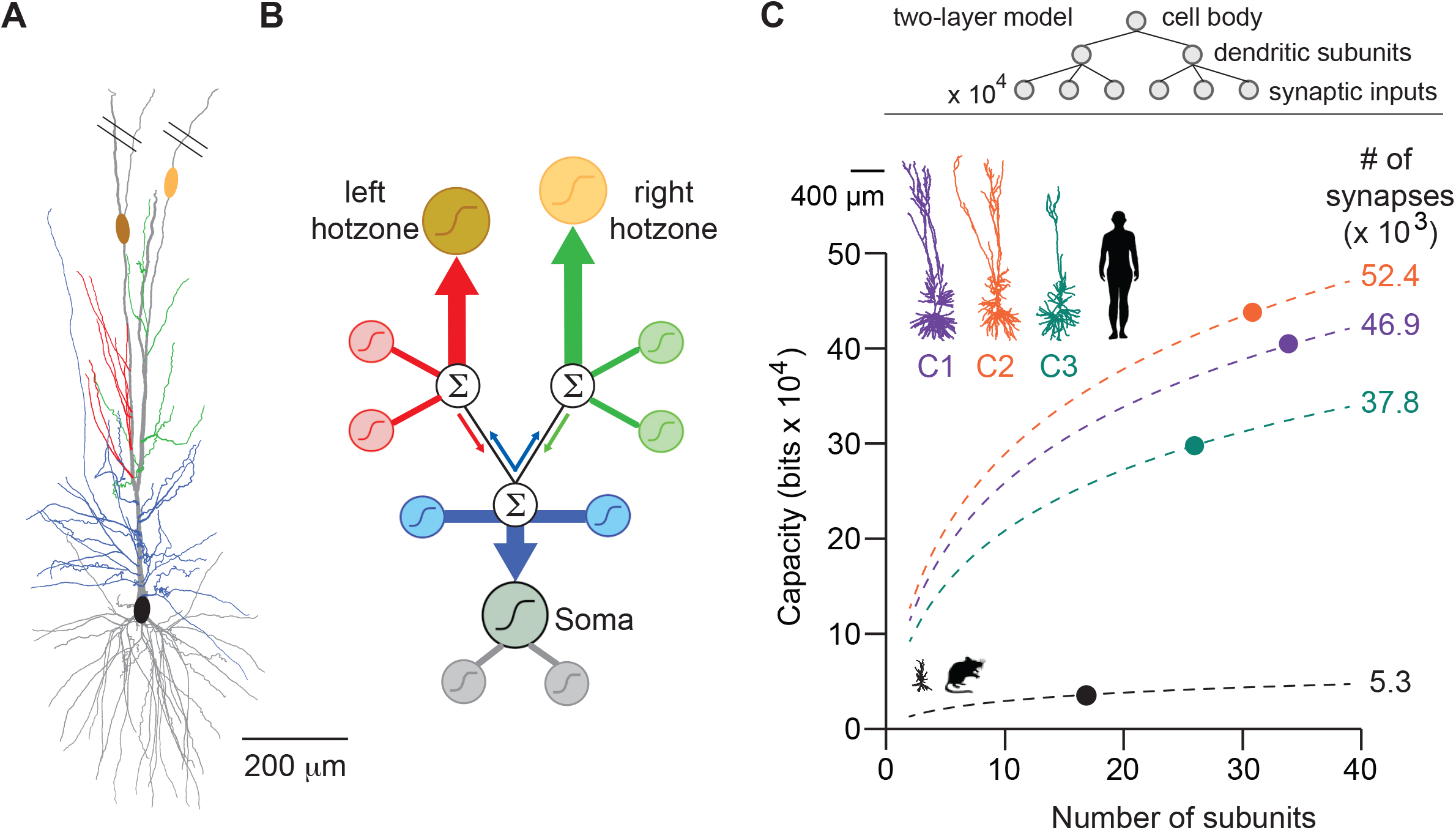
Structural complexity supports computational richness. (A) Example morphology (same as Fig.4-6) at increased magnification. (B) Cartoon of representative morphology illustrating early apical trunk bifurcation that gives rise to independent nonlinearities (NMDA spikes) in individual oblique dendrites (green, red, and blue sigmoidal thumbnails) and preferential routing of these NMDA spikes to different parts of the dendritic tree (thick and thin color-coded arrows). These local nonlinearities, combined with preferential signal routing and interactions between different dendritic compartments (summation symbol), ultimately gives rise to complex nonlinear input/output transformation in human CA1 pyramidal neurons. (C) Computational capacity differs for the three main classes of human CA1 pyramidal neurons when considered as a two-layer model; the capacity depends on the number of non-linear dendritic subunits per neuron and the total number of synapses. The average number of contacts per connection was assumed to be five in both cases (d = s/5 for the parameters used in ^80^. The thick dot on each line illustrates the average multi-layered model for the different species and cell classes. Note: computational capacity of human CA1 pyramidal neurons is 8 to 12-fold higher compared to mouse due to branching characteristics, and increased numbers of dendritic subunits and spine count.

To conclude, we quantified the morpho-electric and computational properties of three types of human hippocampal CA1 pyramidal neurons. This revealed the preferential channeling of information to apical hot zones or soma, the impact of non-linear Ca^2+^ transients in the dendrites on somatic Na^+^ spikes and large number of independent subunits which consistently endows human *CA1* pyramidal neurons with powerful computational capabilities.

## Discussion

The hippocampal formation is responsible for evolutionary conserved behaviors such as spatial navigation, learning, and memory encoding/consolidation. These cognitive functions rely on passive and active biophysical properties of pyramidal neurons that were never characterized in human hippocampus. Here, we used acute resection samples of non-pathological human hippocampus and uncovered previously unknown structural and biophysical characteristics of these cells. We show that i) h*CA1* pyramidal neurons have extended dendritic architecture and many oblique dendrites (Fig.1), ii) neurons consistently show resonant properties and the preferred frequency (Fig.2) corresponds to the lower theta frequencies recorded in human subjects, iii) multi-variable cluster analysis reveals three main classes with divergent morpho-electric properties (Fig.3) and iv) morpho-electric properties lead to non-typical routing of dendritic activity (Fig.5), dendritic nonlinearities and increased computational capabilities (Fig.6, 7). Thus, h*CA1* pyramidal neurons differ from mouse *CA1* pyramidal neurons in all properties studied, including increased structural complexity and enriched computational capacity.

The resonance frequency of h*CA1* pyramidal neurons (2.9 Hz) we uncovered in single neuron recordings closely matches the in vivo theta rhythm of human hippocampus (i.e. 1-4 Hz, ^17,19,57^. The temporal domain at which human hippocampus operates is thus slower compared to rodents as theta-rhythm in mice and rats occur at higher frequencies (i.e. 4 - 10 Hz, ^2,20,58^. We found that the preferred frequency of human (2.9 Hz) and mouse (2.9 Hz) *CA1* pyramidal neurons are highly comparable (Fig.2, Suppl.Fig.2). This indicates that theta rhythm in mice (4 – 10 Hz) and preferred frequency of mouse *CA1* pyramidal neurons (2.9 Hz) do not match. In contrast, theta in human (1-4 Hz) and preferred frequency in human *CA1* pyramidal neurons (2.9 Hz) overlap. Furthermore, while nearly all neurons had resonant properties in humans, a large fraction of neurons was non-resonant in mice. Our computational modeling (Fig.4-6) showed the importance of I_h_. First, I_h_ increases somatic excitability and adds nonlinear properties to the dendritic tree. Second, HCN channel activity results in dendrites that are sensitive to the frequency of synaptic inputs, and this property could be pivotal for phase-locking during spatial navigation ^2,59^ or combining neuronal assemblies during working memory ^12,22^.

In rodent *CA1* pyramidal neurons, it was shown that synaptic activity in individual dendritic branches triggers local NMDA spikes ^41,42^. This could support branch-selective integration of synaptic inputs ^33,34^ or branch-constrained synaptic plasticity ^60^. When multiple dendritic branches are activated simultaneously, the more global depolarization can be sufficient to activate dendritic voltage-dependent calcium channels (ie. trigger a Ca^2+^ spike), resulting in high-frequency bursts at the soma^42,55,61,62^. We show that the dendritic architecture of human *CA1* pyramidal neurons facilitates such compartmentalization, nonlinearities and computational complexity. First, apical dendrites in 69% of our reconstructions show at least one early bifurcation in the apical trunk (Fig.3, see also ^37^ versus mouse 40%) and second, we show the abundance of oblique dendrites on the apical dendritic tree. Our simulations reveal the impact of this particular dendritic architecture (Fig.7): electrical signals do not effectively travel to neighboring oblique dendrites and thus act as independent compartments (Suppl.Fig.4). This is highly relevant for both information encoding and branch-specific plasticity. We show preferential routing of electrical signals in specific parts of the apical dendrite and finally, we show that multi-site NMDA spikes can trigger dendritic Ca^2+^, somatic Na^+^ spikes or both. In summary, the combination of extensive morphologies, compartmentalization of dendrites and different forms of nonlinearities results in a computational unit with complex input-output function ^63,64^.

Our measurements of resonant properties and computational complexity bridge theories on memory function of human hippocampus and may provide a mechanistic explanation on the single-cell level that ultimately extrapolates to human-specific memory capacity and mental flexibility. Average resonance frequency was also remarkably consistent across subjects. This may be surprising considering divergent demographics for surgical cases such as age, genetic background, or disease history, but could also imply this feature is a hard-wired, fundamental intrinsic property of h*CA1* pyramidal neurons. The cautionary note is however that we cannot exclude the possibility that morpho-electric properties were affected by disease history or anti-epileptic medication, even in the absence of structural alterations in our resection tissue. Similarly, it is impossible to determine true biological variability in our data or variability that emerged due to medical history. It is promising though that larger datasets on human resection tissue failed to reveal a link between disease history and dendritic architecture, sag current or related structural/functional properties ^65–67^.

We also found that our human *CA1* pyramidal neurons can be classified into three main classes with divergent morpho-electric properties that are also apparent during computational simulations (Fig.3-7). This is important in view of cellular diversity in rodent *CA1* which has several axes, including transcriptional profile ^68^, preferred theta phase ^69^, spiking properties ^70,71^, strength of action potential backpropagation ^72^ or neuronal properties associated to dorsal/ventral soma location ^27^. Based on genetic profiling in human hippocampus, two major classes of pyramidal cell-types were put forward in hippocampus ^73,74^. Alternatively, apical trunk morphology was used to subcategorize morphologies ^37^, but in general, cellular diversity remains largely unexplored for h*CA1* pyramidal neurons. In rodent *CA1*, cellular diversity maps onto sublayers of rodent *CA1* as pyramidal neurons located in superficial and deep sublayers of SP exhibit clear differences in spiking properties, theta modulation or place field formation ^75^. This raises the question whether the organizational principle of h*CA1* is similar and this topic becomes even more relevant in view of the increased thickness of human *Stratum Pyramidale (SP)* relative to rodents (i.e. rat: 5 cells ^76^ mouse: 75 μm ^37^ or macaque: 10-15 cells ^31^). In our surgical resection samples, we measured *SP* thickness of 1111 μm, which is very similar to the 1130 micrometer reported for brain samples after postmortem donation by healthy subjects ^77^. We prioritized the characterization of resonance properties of neurons as this biophysical property is fundamental to hippocampal function. We carried out our measurements from h*CA1* pyramidal neurons in a relatively central band of the *SP* layer, which was at the expense of sampling across the full width of *SP*. Thus, one important future objective could be to determine whether sublayers or pyramidal sub-categorizations exist in h*CA1*. If so, the next question is whether diversity in *SP* location and additional variability in cellular properties impact resonance frequency.

Across species, *CA1* pyramidal neurons consistently show a large number of oblique dendrites ^30,31,45^. These oblique dendrites show highly specialized properties including integrative properties ^32–34^, excitability ^41^, and signal propagation ^32,78^. Here, we show that the particularly large number of oblique dendrites and the resultant large membrane area could impose a severe “on path” conductance load which might severely decouple the soma from the tuft dendrite. However, this effect is partly compensated for by the early (proximal) bifurcation of the apical tree and the respective split of the oblique dendrites. Consequently, some obliques arise from the proximal (pre-branch) apical dendrites and the rest of the obliques split between the two apical branches. We show computationally that this specific dendritic cable design still allows for soma-to-tuft communication and at the same time, allows preferential routing of information from the proximal obliques to the soma and from the two distal obliques’ subgroups to their respective apical branch where nonlinear electrogenesis mechanisms are likely to operate. Furthermore, this splitting of the obliques into three subgroups, due to the early branching of the apical tree, enables these subgroups of oblique dendrites to operate as independent functional subunits. We also demonstrated the frequency-dependency of this preferential communication, which results from asymmetrical cable structure of h*CA1* neurons around the proximal apical bifurcation. This frequency-dependency may further enhance the computational capabilities of these neurons e.g., for coding multiple locations in space (multiple place fields). Since individual oblique dendrites may represent an additional site for action potential or NMDA spike generation ^79^, the large number of oblique dendrites in h*CA1* pyramidal neurons boosts computational (and memory) capacity of these neurons ^33,64,80^. Such computational capabilities are expected to be dramatically lower for rodent CA1 pyramidal neurons, since the architecture of the apical dendrite does not show similar topology, resulting in a more compact dendritic cable ^37^.

To conclude, we show that h*CA1* pyramidal neurons have elaborate dendritic trees and morpho-electric properties that support many local nonlinear operations during input-output transformations. This translates to enriched computational complexity and thus encoding and memory capabilities. h*CA1* pyramidal neurons show a clear frequency preference which is consistent across neurons and subjects. The ability of h*CA1* pyramidal neurons to respond to a preferred frequency causally depends on HCN channel function ^81^ and the preferred frequency accurately matches the hippocampal theta rhythm observed during complex human behaviors. The combination of deep brain recording techniques ^82,83^ and single cell recordings in non-sclerotic human resection tissue for transcriptomic classification of cell-types ^66^ paves an exciting way for uncovering additional unknown aspects of human hippocampus function including genes, cells, circuits and ultimately cognitive behavior.

## Lead contact

Further information and requests for resources and reagents should be directed to and will be fulfilled by the lead contact, Christiaan de Kock.

## Materials availability

This study did not generate new unique reagents.

## Data and code availability

All data from the figures is available at the DataverseNL link: to be added

Morphological reconstructions, raw electrophysiology or modeling data reported in this paper will be shared by the lead contact upon request. Analysis code has been deposited at Github and is publicly available as of the date of publication, as seen in the key resource table. Any additional information required to reanalyze the data reported in this paper is available from the Lead Contract upon request.

## EXPERIMENTAL MODEL AND SUBJECT DETAILS

### Human surgical specimens

All procedures were performed with the approval of the Medical Ethical Committee of the VU medical center (VUmc), and in accordance with Dutch license procedures and the Declaration of Helsinki. All patients provided written informed consent.

Data included in this study were exclusively obtained from (non-pathological) neurosurgical tissue resections for the treatment of temporal lobe epilepsy (n = 4) or epilepsy with tumor (n = 1) or unknown treatment (n=1) in 4 male and 2 female patients (Table 1). We did not detect any influence of gender on the morphological/physiological properties.

### Mouse specimens

All procedures involving mice were approved by the animal ethical care committee of the VU university. Mixed strains of male (n=2) and female (n=2) mice from 63-70 days old were used for experiments. Mice were maintained on a 12 h light/dark cycle in a temperature and humidity-controlled room. Mice were housed 3-6 per cage with free access to food and water. We did not detect any influence of gender on the morphological/physiological properties.

## METHOD DETAILS

### Human acute brain slice preparation

During surgical treatment of underlying brain pathology, the hippocampus was taken out “en block”, in addition to (partial) resection of the temporal lobe. Structural integrity of resected hippocampus was assessed with 1) presurgical MRI, 2) assessment of the cytoarchitectural integrity by an expert pathologist, 3) differential interference contrast images during electrophysiology, and finally 4) post-hoc histology of tissue used for electrophysiology (NeuN and biocytin-DAB). We obtained hippocampal tissue from a total of 6 patients: 5 patients from the VU Medical Center (VUmc) and 1 patient from Harborview Medical Center (Table 1). In these hippocampal specimens, presurgical MRI and posthoc histology did not reveal structural abnormalities (Table 1).

Immediately upon resection, the tissue block containing hippocampus proper was placed into a sealed container filled with ice-cold, artificial cerebral spinal fluid (aCSF). For two samples, initial aCSF consisted of (in mM): 110 choline chloride, 26 NaHCO_3_, 10 D-glucose, 11.6 sodium ascorbate, 7 MgCl_2_, 3.1 sodium pyruvate, 2.5 KCl, 1.25 NaH_2_PO_4_ and 0.5 CaCl_2_. For five six additional samples, aCSF consisted of (in mM): 92 *N*-methyl-d-glucamine chloride (NMDG-Cl), 2.5 KCl, 1.2 NaH_2_PO_4_, 30 NaHCO_3_, 20 4-(2-hydroxyethyl)-1-piperazineethanesulfonic acid (HEPES), 25 d-glucose, 2 thio-urea, 5 sodium-l-ascorbate, 3 sodium pyruvate, 0.5 CaCl_2_.4H_2_O and 10 MgSO_4_.7H_2_O. Before use, the aCSF solution was carbogenated with 95% O_2_, 5% CO_2_ and the pH was adjusted to 7.3 by addition of 5M HCl solution, after which it was put on ice until use.

The transition time between resection in the operation room and arriving at the neurophysiology lab was < 20 minutes. Immediately upon arrival at the lab, slice preparation commenced. First, residual blood was rinsed from the tissue block, while remaining submerged in ice-cold NMDG solution. Next, orientation of the tissue for slicing was determined based on gross macroscopic anatomical hallmarks of the hippocampus proper and adjacent structures.

Next, the tissue block was glued onto the slicing platform such that the slice angle ensured optimally intact (apical) dendrites, and 350-μm-thick coronal slices of the hippocampal body were prepared using a vibratome (Leica V1200S), in ice-cold aCSF solution. Each slice was then transferred to a warmed holding chamber (34 °C), containing NMDG-based aCSF, for 12 minutes under constant carbogenation. Next, slices were transferred to a holding chamber with aCSF containing (in mM): 92 NaCl, 2.5 KCl, 1.2 NaH_2_PO4, 30 NaHCO_3_, 20 HEPES, 25 d-glucose, 2 thiourea, 5 sodium-L-ascorbate, 3 sodium pyruvate, 2 CaCl_2_.4H_2_O and 2 MgSO_4_.7H_2_O (pH 7.3) and stored at room temperature for at least an hour until used. The time between tissue resection and electrophysiological recordings could thus vary between 2 - 15 hours. Osmolarity of various aCSF solutions were set to 310 mOsm, using the Vapro (5600 Vapor pressure, Elitech) osmometer.

### Mouse acute brain slice preparation

All procedures related to mice were approved by the animal ethical care committee of the VU university. C57BL/6 mice (n = 4, age 63-70 days, 2 males, 2 females) were anaesthetized with euthasol (i.p., 120 mg/kg in 0.9% NaCl), and transcardially perfused with 10 mL ice-cold carbogen-saturated NMDG solution. Upon removal of the brain, 350 μm thick coronal slices were obtained as described above.

### Electrophysiology

Recordings were obtained at 34°C, in aCSF containing (in mM): 125 NaCl, 3 KCl, 1.2 NaH_2_PO_4_, 1 MgSO_4_, 2 CaCl_2_, 26 NaHCO_3_, and 10 d-glucose (310 mOsm), continuously bubbled with 95% O_2_, 5% CO_2_. In a subset of experiments, recordings were made in aCSF containing (in mM): 126 NaCl, 2.5 KCl, 1.25 NaH2PO4, 26 NaHCO3, 12.5 glucose, 2 CaCl2·4H2O, 1 MgSO4·7H2O, 1 Kynurenic acid and 0.1 picrotoxin (pH 7.3). Borosilicate glass patch pipettes (3-6 MΩ resistance) (Harvard apparatus/Science products GmbH) were filled with intracellular solution, containing (in mM): 115 K-gluconate, 10 HEPES, 4 KCl, 4 Mg-ATP, 10 K-Phosphocreatine, 0.3 GTP, 0.2 EGTA, and biocytin 5 mg/ml, pH adjusted to 7.3 with KOH, osmolarity 295 mOsm/kg). For patch-seq experiments, internal solution contained (in mM): 110.0 K-gluconate, 10.0 HEPES, 0.2 EGTA, 4 KCl, 0.3 Na2-GTP, 10 phosphocreatine disodium salt hydrate, 1 Mg-ATP, 20 μg/ml glycogen, 0.5U/μL RNAse inhibitor (Takara, 2313A), 0.5% biocytin and 0.02 Alexa 488. The pH was adjusted to 7.3 with KOH, osmolarity to 295 mOsm/kg.

Whole-cell patch-clamp recordings were made with a Multi-Clamp 700B (Molecular Devices), digitized at a rate of 20 kHz (ITC-18 computer interface instrutech corporation / National Instruments USB-6343) and obtained with either PClamp 10 (Molecular Devices) or MIES (Allen institute) in IgorPro 8 (WaveMetrics). Files were stored in Axon Binary File (abf) and Neurodata Without Borders (nwb) format, respectively. Prior to recording the pipette offset was compensated for, and in current clamp the bridge was balanced. Pipette access resistance was monitored throughout the recording, and was between 5-18 MΩ. Recordings with a pipette access resistance >20 MΩ were excluded from the dataset. HCN channel antagonist ZD7288 (10 μM; Hello bio) or voltage-gated calcium channel blocker cadmium (CdCl_2_) (10 μM; J.T. Baker) were prepared in recording aCSF, and washed in upon completion of all protocols, after which the protocols were repeated.

### Immunohistochemistry

After recordings, slices were fixed in 4% paraformaldehyde (in phosphate buffer) for a minimum of two days. Subsequently, biocytin-filled neurons were recovered using the chromogen 3,3-diaminobenzidine (DAB) tetrahydrochloride avidin–biotin–peroxidase method. Both at the department of (Neuro)Pathology at the UMC and our lab, the neurotypical status of the tissue was evaluated using a staining for Neuronal Nuclei (NeuN) using rabbit-anti-NeuN primary antibody (ThermoFisher, catalog number PA578499) and Biotinylated Affinity Purified Goat Anti-Rabbit IgG secondary antibody (ThermoFisher, 32054) for a subset of tissue samples. Slices were mounted on slides and embedded in mowiol (Clariant) or Aqua-Poly/Mount (Polysciences).

## QUANTIFICATION AND STATISTICAL ANALYSIS

### Neurophysiology

We applied multiple stimulation protocols to obtain passive and active (sub- and suprathreshold) properties of the human *CA1* pyramidal neurons. All protocols started at an imposed membrane potential of -70 mV ± 1 mV.

#### Passive properties

To assess passive subthreshold properties, hyperpolarizing square 1s current injections were applied, starting from -150 pA, with increments of +25 pA. A -25 pA square current injection was used to calculate the input resistance and membrane time constant (tau). HCN channel activity was quantified on sweeps with current injections leading to 3 mV hyperpolarization (to compensate for the difference in response between recordings due to input resistance variability). The difference between sag peak and steady-state hyperpolarizing responses was defined as delta sag peak (in mV). To generate the sag ratio, the delta sag peak was divided by the steady state hyperpolarization response of the same sweep.

Next, resonance frequency (Fig.2) was determined from the voltage response (± 5-10 mV) to a sinusoidal current injection with a frequency range of 0.5 to 30 or 40 Hz (in 10 or 20 s, respectively). Next, the impedance amplitude profile (ZAP) was derived via the ratio of the fast Fourier transform of the voltage response to the fast Fourier transform of the sinusoidal current injection. The resonance frequency (F_res_) reflects the frequency at which maximum impedance was observed.

#### Active properties

To obtain active, suprathreshold membrane properties, a 1s depolarizing square current injection was applied, ranging from +25 pA to 1000 pA, with increments of 25 pA. The smallest current injection that resulted in an action potential (AP), was set as the rheobase value, and this first sweep exceeding AP threshold was used for all analyses of active suprathreshold membrane properties (AP threshold, AP amplitude, AP upstroke, AP downstroke and AP halfwidth). AP threshold was defined as the point where the slope exceeded 23 mV/ms. AP amplitude was specified as the voltage change between the AP threshold and the AP peak. AP halfwidth was defined as the width of the AP (in ms) at half maximal amplitude. Upstroke and downstroke speed (mV/ms) are computed as the mean rising and fall speed between 30 and 70% of the AP, respectively.

#### After-depolarization properties

We quantified the amplitude of the after-depolarization (ADP) during somatic AP bursts to probe dendritic electrogenesis ^61^. Three short (3 ms) pulses were given at increasing frequencies. The voltage traces were low-pass filtered at 1200 Hz with a second-order Butterworth filter. Only sweeps that successfully triggered 3 APs were included in the analysis. Spiking frequency was computed as: F= 2/(t_(spike 3)-t_(spike 1)) where t is the timestamp of a given spike (at AP peak voltage). All traces were baseline-corrected by subtracting the median membrane voltage. ADP amplitude was measured at 7.25 ms after the peak of the action potential (average ADP peak time). ADP peak was computed as the maximum voltage value after the AP derivative became positive (after AP peak).

For each neuron, the relation between ADP amplitude and spiking frequency was fitted with three different functions: a sigmoid, a saturating increasing exponential and a linear function. The best fit was chosen as the one yielding the maximum R-squared value. For all neurons, the best fit was the exponential fit:

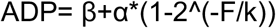

Where ADP is the ADP amplitude, F is the firing frequency, α and β are amplitude and position parameters and k is the half-maximum frequency.

#### Control for burst spiking

Burst spiking was not part of the features as we typically encountered a single spike (26 out of 41 recordings) upon rheobase stimulation or multiple spikes at low frequencies (3.2 Hz, 2.1 - 6.3 Hz, median, 1st - 3rd Quartile range, n = 15). We found only n = 1 pyramidal neuron with instantaneous frequency of 73 Hz at rheobase current injection, but this is still considered regular spiking ^70^. At maximal current injection, only n = 2 neurons exhibit an instantaneous frequency exceeding 100 Hz (interspike interval (ISI) cell 1 = 9.4 ms, ISI cell 2 = 9.5 ms) and could be considered bursting neurons, but only upon maximal current injection. Upon short, 3 ms current injections, human *CA1* pyramidal neurons are capable of spiking at high frequencies (> 100 Hz, see below) and we thus conclude that prolonged somatic current injection does not favor burst spiking in our human resection samples.

### Morphological reconstruction

All recovered neurons underwent critical quality assessment of staining quality and occurrence of slicing artifacts. Only neurons that passed quality control were reconstructed with Neurolucida software (Microbrightfield, Williston, VT, USA), using an 100x oil objective. Using DAB and NeuN histology, *Stratum Pyramidale* (*SP*) could be unambiguously identified, as well as the border between *SP* and *Stratum Radiale* (*SR*). The border between *SR* and *Stratum Lacunosum Moleculare* (*SLM*) was estimated based on differences in contrast for *SR* and *SLM* layers using light microscopy. Finally, reconstructed *CA1* neurons were annotated relative to layer borders.

### Statistical analysis

Data were analyzed using analysis scripts written in Matlab 2021A (MathWorks) and statistical analyses were performed using Prism 7.2 (GraphPad Software). All AP analyses were performed using customized Matlab scripts (Source code available at https://github.com/ElineJasmijn/Morphys, taken and adjusted from https://github.com/INF-Rene/Morphys). Non-parametric data are visualized in boxplots (generated in Matlab 2021a) with the central mark as the median, the edges of the box the 25th and 75th percentiles, the whiskers extending to the most extreme data points, excluding the outliers. In all boxplots, each dot represents a single morphology or recording.

### Cluster analysis

We used 22 structural and biophysical features from n = 31 digital reconstructions and electrophysiological recordings and this subset with bimodal data originated from 3 patients. To reduce the dimensionality of the dataset and identify the most informative features for clustering, we performed a principal component analysis (PCA) ^85^. Only principal components that explained more than 5% of the variance were included in the dataset, resulting in a total of 7 principal components, which together explained approximately 83.6% of the variance. We next used unsupervised hierarchical clustering and clustering results were visualized using a 2-dimensional dendrogram and t-SNE plot (Fig.3).

### Modeling

#### Extracting passive cable and HCN channel properties

We constructed detailed compartmental models for 13 human *CA1* pyramidal neurons that were both 3D reconstructed and physiologically characterized. These models included passive membrane properties (R_m_, C_m_, R_a_) extracted by fitting the models to the experimental results (Fig. 4). These models also incorporated HCN channels that were distributed over the dendritic surface (see below). In order to compensate for the absence of spines (and thus underestimating true membrane surface area) in our morphological reconstructions, we used the “F factor” method to globally integrate these spines into the dendritic membrane, using spine parameters shown in Table S1 ^86^. Models were fitted to experiments by minimizing the root-mean-square-deviation (RMSD) between the somatic voltage response of the model and the respective experimental voltage response, for both transient voltage in response to a short (2 ms) hyperpolarizing current injection (not shown), as well as the I_h_ related sag’s half width, sag’s amplitude, minimal voltage response and resting membrane potential responses to several long (1 s) hyperpolarizing step currents (see Fig.4, i.e. training sweeps). For these fits we used the genetic algorithm developed under BluePyOpt ^84^, on 100 individuals for 1000 generations (the parameters bound for the fits are listed in Table S2). In order to evaluate the quality of fit, we tested the model on the previously unused 1s long pulses.

#### Simulating excitatory synapses

Excitatory synaptic inputs impinging on the dendritic tips were modeled as composed of both AMPA and NMDA receptors.

The synaptic current, *I*_syn_, was modeled as:

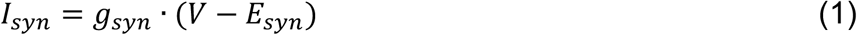

where *g*_*syn*_ is the transient synaptic conductance and *E*_*syn*_ was set to 0 mV. for both AMPA and NMDA we modeled *g*_*syn*_ as a two-state kinetic synapse:

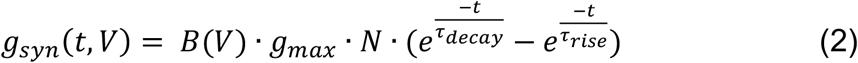

with *g*_*max*_ is the maximal synaptic conductance (for AMPA / NMDA 0.4, 1.2 nS respectively), in case only AMPA synapses where activated g_max_ AMPA was set to 1.6), B – voltage-dependent component and N - normalization factor following Eqs. (3-5):

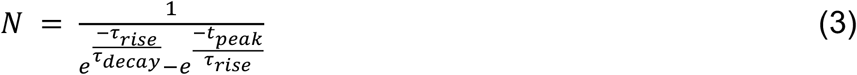

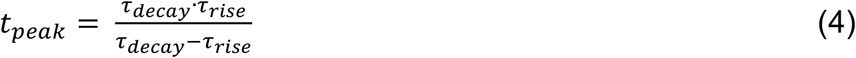

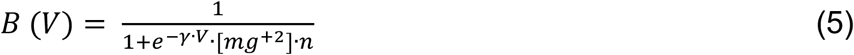

With, τ_rise-AMPA_=0.3 ms, τ_decay-AMPA_=1.5 ms, τ_rise-NMDA_=8 ms, τ_decay-NMDA_=53 ms, γ=0.076 mV and n=0.27 nm.

#### NMDA spike simulation

In order to generate an NMDA spike, we activated 30 synapses per oblique branch with all activated synapses being clustered in the distal segment of the respective tip. In cases where the number of NMDA spikes was larger than the number of dendritic tips (Fig. 6), an additional NMDA spike was generated in one of the terminal branches at a random location.

#### Signal transfer in CA1 dendrites

The key focus of the theoretical part of this work is to explore possible functional implications of the large number, and the specific spatial distribution, of the oblique dendrites in human *CA1*. Towards this end, we have developed a way to graphically depict the “equivalent cable” as seen from different viewpoints; e.g., from the soma view point or from one of the obliques viewpoints (see below).

Our cable models are constructed so that the somatic direction is pointed downwards and the tuft direction is upwards. When the cable is constructed from the somatic viewpoint, the respective equivalent cable results in a basal tree pointing downwards and apical cable upwards (Fig. 4). In cases where the equivalent cable is constructed from an oblique viewpoint, as in the insets of Fig. 5A, the input oblique is marked by a yellow circle (i.e., apical zone) and the equivalent cable of the respective apical branch from which this oblique emerges from points upwards and the rest of the tree points downwards.

This presentation provides a direct intuitive understanding of the impedance load imposed on signal transfer in *CA1* dendrites e.g., by the tree on a specific subgroup of the oblique dendrites. For this graphical representation we computed the “equivalent cable” of *CA1* dendrites (Eq. (6) below) subdivided into the different subtrees composing *CA1* dendrites). X = *x*/λ, *x* is the physical path distance of a given dendritic location from the soma, 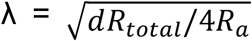 where d is the diameter of a given dendritic branch, *R*_*total*_= 1/(*g*_*leak*_+ *g*_1*h−rest*_). _1*h−rest*_ is the conductance of I_h_ at resting membrane potential.

In Figure 5A, insets, we constructed “equivalent cable” for the respective *CA1* neurons, based on Rall’s cable theory ^87^. The variable diameter, *d*_*eq*_(*X*), of this cable is,

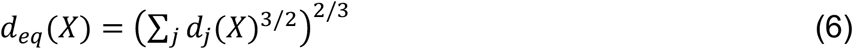

where X is the cable (electrotonic) distance from the point of interest (e.g., the soma or a particular oblique dendrite) and *d*_j_(*X*) is the diameter of the jth dendrite at the distance X from that point of interest. Figure 5A show such equivalent cables as seen from “oblique-centric” perspective; this enables one to graphically appreciate the conductance load (current sink) imposed by different types of obliques (proximal, distal left and distal right obliques) on signal transfer in human *CA1* neurons (see below).

One way to quantify signal transfer from the oblique dendrites to the rest of the dendritic tree is to measure the transfer Impedance (Z_tr_) from any given oblique to other points of interest in the dendritic tree,

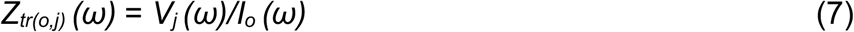

where I_o_ (ω) is the peak current injected to a particular oblique (o) dendrite and V_j_ (ω) is the peak voltage recorded at location j (j = soma, left hotzone, right hotzone) and ω is the input frequency. Note that transfer resistance is the case were ω = 0.

In the present study, we selected three target points of interest: the soma and two putative “hotzones” in the apical tree, and origin points: all oblique tips. These hotzones were manually selected per *CA1* neuron; one at the left apical dendrite distal to the main apical branch (the nexus) and one at the right apical branch (see Fig. 4-6 A,B). Z_tr_ was computed with incorporation of the conductance of the HCN channels at resting membrane potential.

#### Modeling HCN channels

One of the most well-established fingerprints of hippocampal *CA1* pyramidal neurons in rodents is their strong sag voltage, which was shown to result from the presence of dendritic HCN channels. I_h_ current was shown to have important computational implications for synaptic integration for frequency tuning ^28,81,88^. In human supragranular pyramidal neurons, I_h_ was found to be even more pronounced near the resting membrane potential with respect to rodents, due to a leftward shift in its voltage activation ^89^. This was also the case in our models of human *CA1* neurons (see Results). The full set of parameters obtained is listed in Table S3.

I_h_ current, due to the activation of HCN channels, was modeled with a shifted parameter for the voltage activation (as in ^28,88^, as follows:

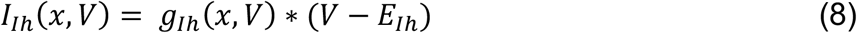

where E_Ih_, the reversal potential for the HCN channels, was set to -45 mV. We distributed HCN as linearly increasing from the soma out, following ^28^:

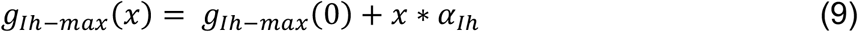

where x is the distance from the soma in μm and α_Ih_ and g_Ih-max_(0) are free parameters for the searching algorithm. We modeled g_Ih_ using a two-state Hodgkin–Huxley like channel (see ^90^:

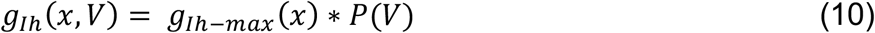

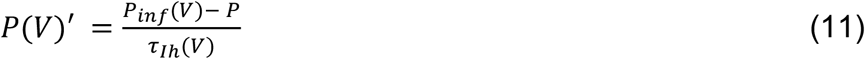

Where P is the probability of the channel to be open, P_inf_ steady state probability and τ_*Ih*_is the time constant of Ih. We allowed a voltage shift in order to compensate for junction potential, V_shift-Ih_, and added a scaling for τ _Ih_, τ _Ih – mul_ and τ _Ih – add_:

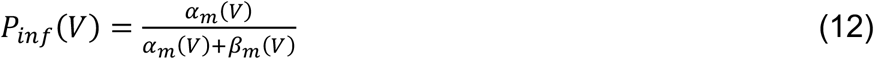

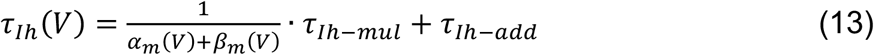

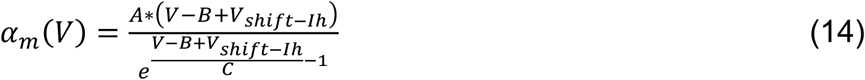

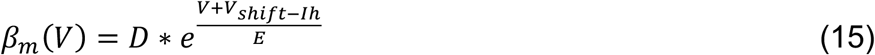

With A = 6.43 s^−1^, B = 154.9 mV, C = 11.9 mV, D = 193 s^−1^ and E = 33.1 mV ^90^.

τ _Ih – mul_, τ_Ih – add_ and V_shift-Ih_ were set to be free parameters for the searching algorithm. This resulted in five Ih-related free parameters for fitting (α_Ih_, g_Ih-max_ at the soma, τ_Ih – mul,_ τ_Ih – add_ and V_shift-Ih_).

#### Modeling Ca^2+^ hotzones and active soma

Voltage dependent Ca^2+^ channels in the hotzones of the apical tree and voltage-dependent Na^+^ and K^+^ channels in the soma were modeled using Hodgkin & Huxley formalism as follows:

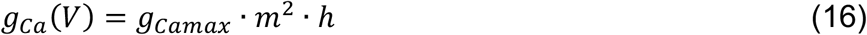

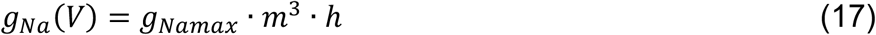

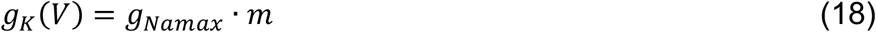

There are two types of voltage-dependent calcium channel conductances: 1) the high-voltage dependent Ca^2+^ conductance with *g*_*Camaxhigh*_ = 10 pS/μm^2^ and 2) a low-voltage dependent Ca^2+^ conductance with *g*_*Camaxlow*_= 800 pS/μm^2^. The somatic voltage dependent Na^+^ conductance *g*_*Namax*_ = 7000 pS/μm^2^ and the voltage dependent K^+^ conductance *g*_kmax_ = 1750 pS/μm^2^.

The activation function *m* (as well as that of the inactivation, *h*) is described as follows:

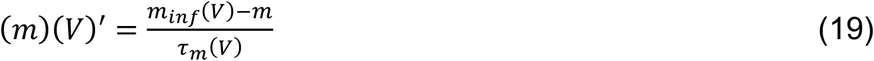

And similarly for *h* for the low-voltage dependent Ca^2+^ channels:

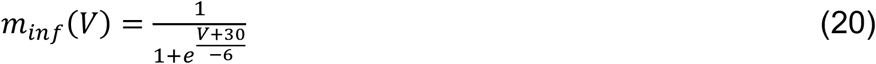

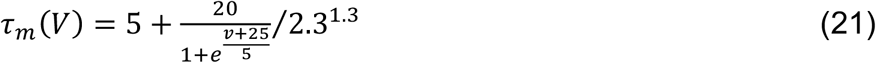

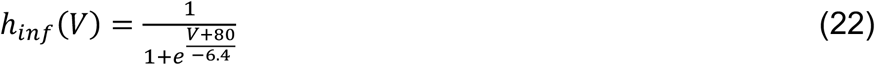

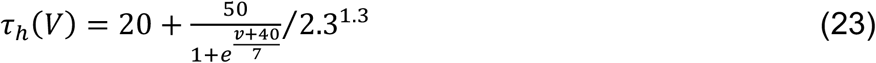

for the high-voltage dependent Ca^2+^, Na^+^ and K^+^ conductances, *m*_*inf*_ and *τ*_*m*_ (and similarly for *h*) were computed as follows:

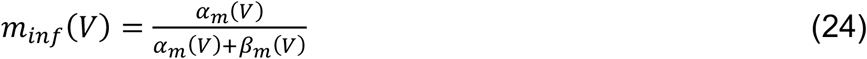

with the exception for:

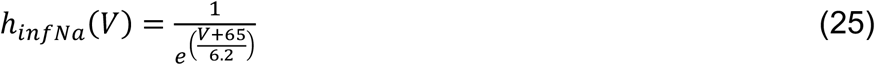

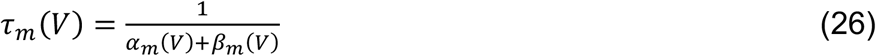

and:

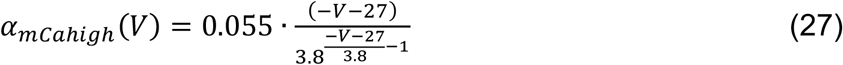

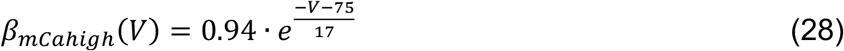

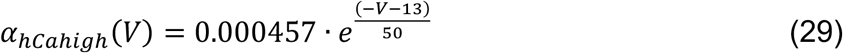

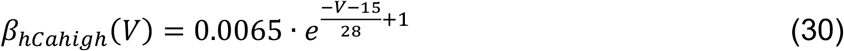

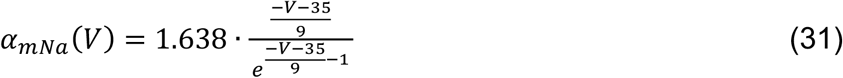

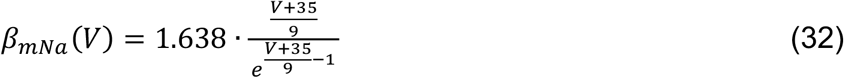

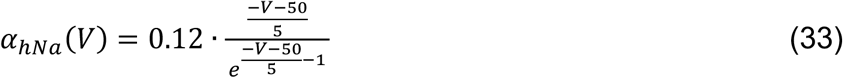

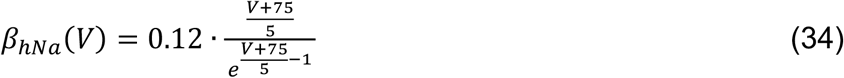

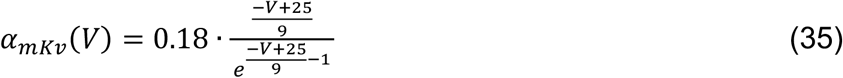

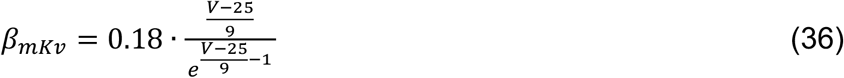

#### Calculation of memory capacity

Memory capacity of hCA1 neurons was computed using the formula ^80^:

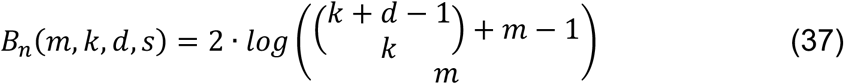

where *m* is the number of dendritic subunits, *s* is the total number of synapses, *d* is the number of NMDA-spikes needed for generating a somatic Na^+^ spike and *k* is s/m, the number of synapses in each subunit. Based on our simulation results (Fig.6) we used d = 5 in the computation of memory capacity since 5 NMDA spikes result in 50% chance to elicit an Na^+^ spike.

Computational capacity was determined for the 3 classes of *hCA1* pyramidal neurons separately. Total number of synapses was the product of total dendritic length (Fig.3) and branch-specific spine density ^37^. The number of dendritic subunits was estimated (a lower bound) as the sum of the number of primary basal dendrites emerging from the soma, the number of oblique dendrites emerging from the apical trunk and the number of tufts’ tips. To validate that these dendritic subunits are indeed independent units, we simulated voltage response after activation of single (Suppl.Fig.4E) or multiple (Suppl.Fig.4F) NMDA spikes at different dendritic locations. The peak depolarization (color coded) remained local, indicating that indeed dendritic subunits act as independent units. For mouse we determined total dendritic length of n = 5 representative morphologies from ^37^ and multiplied these values with branch-specific spine density to compute total number of synapses (s = 5312, ^91^). In addition, m = 17 is based on the same n = 5 representative morphologies. Finally, d = 5.

## Author contributions

Conceptualization, BK, JT, ES, HDM, IS, CPJdK

Methodology, EJM, YL, JP, AAG, FW, TSH, MBV

Software, EJM, YL, JP, FW, RW, DH, MBV, BEK

Investigation, EJM, YL, JP, AAG, FW, TSH, RW, DH, NAG, MBV, EA, BL, BK, JT

Formal analysis, EJM, YL, JP, JM

Funding acquisition, BEK, BRL, ESL, JT, HDM, IS, CPJdK

Resources, SI, RPG

Supervision, HDM, IS, CPJdK

Visualization, EJM, YL, JP, FW, HDM, CPJdK

Writing – original draft, EJM, YL, HDM, IS, CPJdK

Writing – review & editing, EJM, YL, HDM, IS, CPJdK, commented by all authors.

## Data and materials availability

All data needed to evaluate the conclusions in the paper are present in the paper and/or the Supplementary Materials. Analysis scripts are available at https://github.com/AllenInstitute/MIES or https://github.com/ElineJasmijn/Morphys.

## Supplementary Materials

Figs. S1 to S5

Tables S1 to S3

**Supplementary Fig. 1:**
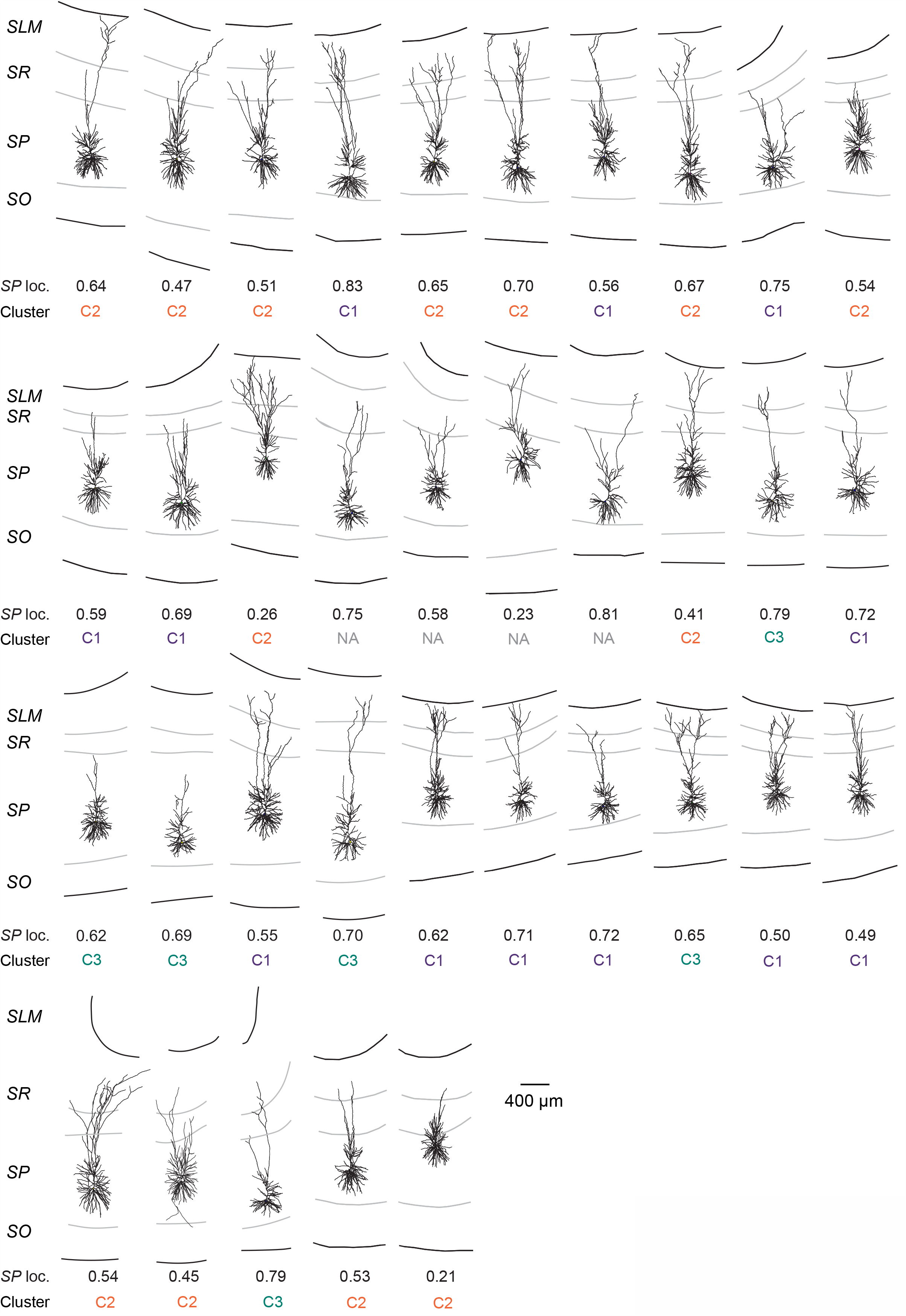
Gallery of n = 35 human CA1 pyramidal neurons used for morphological analysis (Fig.1) and unbiased cluster analysis (subset of n = 31 with morphology <u>and</u> electrophysiology), annotated according to layer position. For the n = 31, each of the electrophysiological features used for cluster analysis was mapped. SP: Stratum Pyramidale, SO: Stratum Oriens, SR: Stratum Radiatum, SLM: Stratum Lacunosum Moleculare. The value “fraction in SP” refers to the position of the cell body within the SP according to the measure “(distance Soma - SLM)/(distance SLM - SO)”. Cluster identity (in color code matching Fig.3) is provided below each reconstruction. NA: not applicable (morphologies not included in cluster analysis).

**Supplementary Fig. 2.**
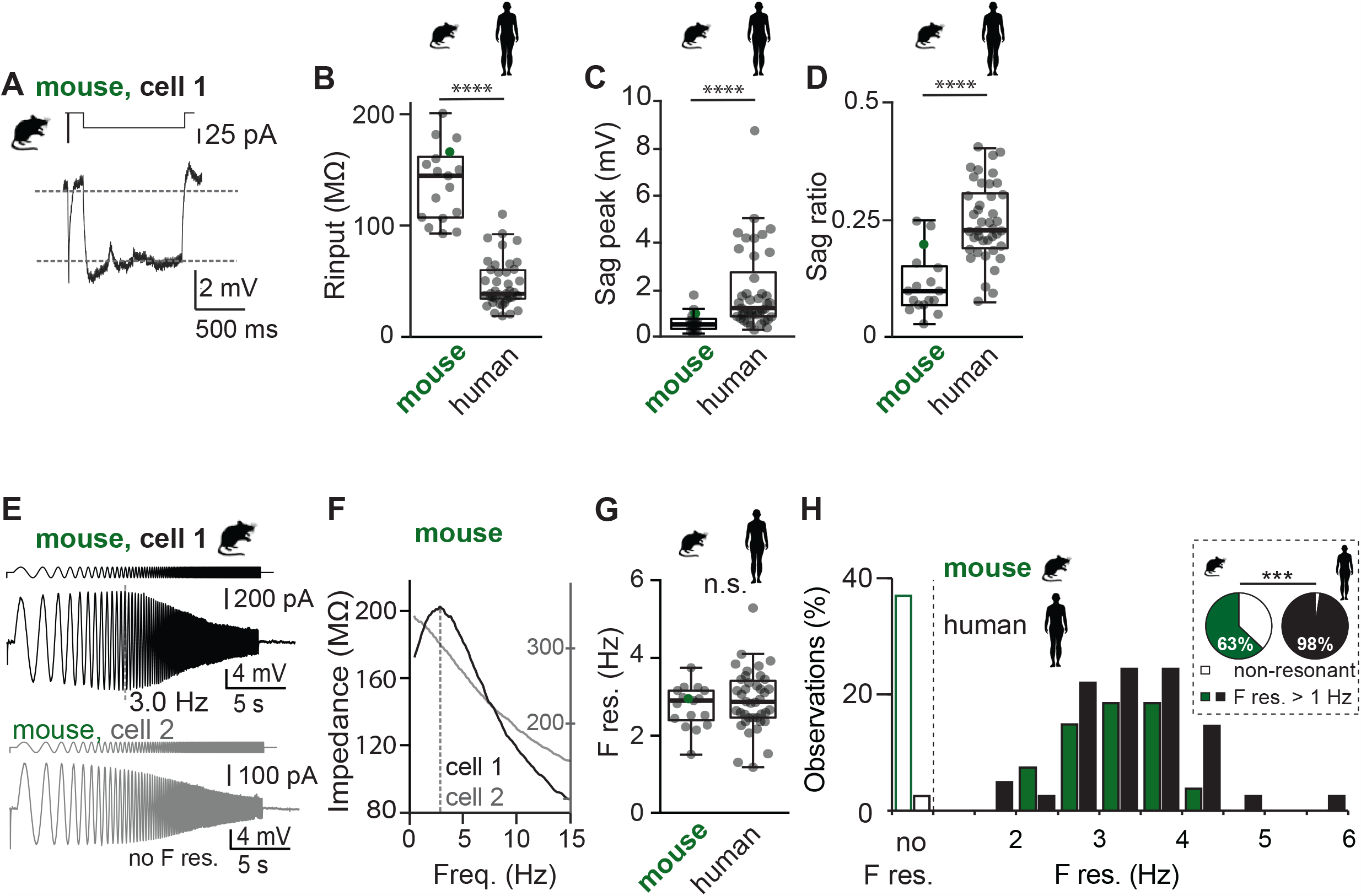
Mouse CA1 neurons resonate at similar frequency, but the incidence is significantly lower. (A) Example mouse CA1 recording with hyperpolarization current to uncover I_h_ (or sag current). Dashed lines correspond to -70 mV and steady-state hyperpolarization (close to -73mV, see Methods), respectively. (B) Input resistance is significantly lower in human compared to mouse CA1 pyramidal neurons. (C, D) Maximal sag current amplitude (C) and sag ratio (D) is significantly higher in human compared to mouse CA1 pyramidal neurons. (E) Chirp protocol to extract resonance properties from mouse CA1 pyramidal neurons. Note two examples, of which cell 2 does not show resonant properties under control conditions. (F) Impedance profiles of example cells 1 and 2. Impedance of cell 2 is illustrated by 2nd y-axis. (G) Population statistics for preferred frequency (i.e. resonance frequency, n = 17, 2.9 Hz, 2.5 - 3.2 Hz, median, 1^st^ – 3^rd^ Quartile). (K) Histogram with distribution of resonance frequencies for human (black) and mouse (green), in addition to fraction of neurons without preferred frequency (open bars). Inset: pie charts representing the fraction of neurons with and without resonant properties. Note that in mouse, a large fraction of recordings did not show preferred resonance frequency (Human: 2.4% vs mouse: 37.0%, Fisher exact test, p < 0.001).

**Supplementary Fig. 3.**
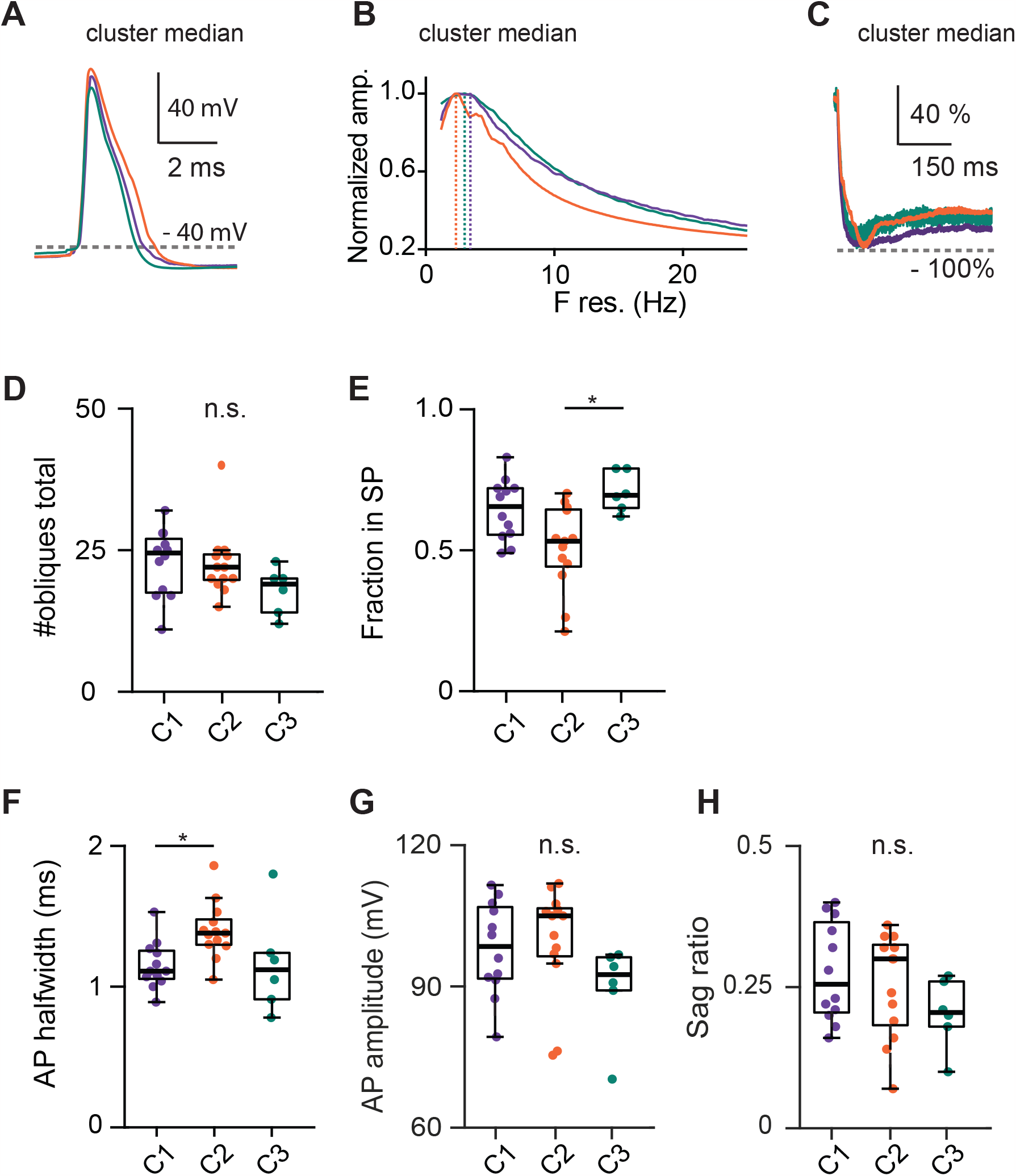
Human CA1 contains three types of pyramidal neurons with distinct morpho-electric properties. (A-C): Population medians for AP waveform (A), resonance properties (B) and sag current upon hyperpolarizing current injection (C), respectively. (D-E) Population statistics for morphological (# of obliques total, D) and anatomical features (Fraction in SP, E) of the three main cell classes. (F-H) Population statistics for electrophysiological features of the three main cell classes. Statistics: Kruskal-Wallis with Dunn’s post-hoc test, * p < 0.05, n.s.: not significant.

**Supplementary Fig. 4.**
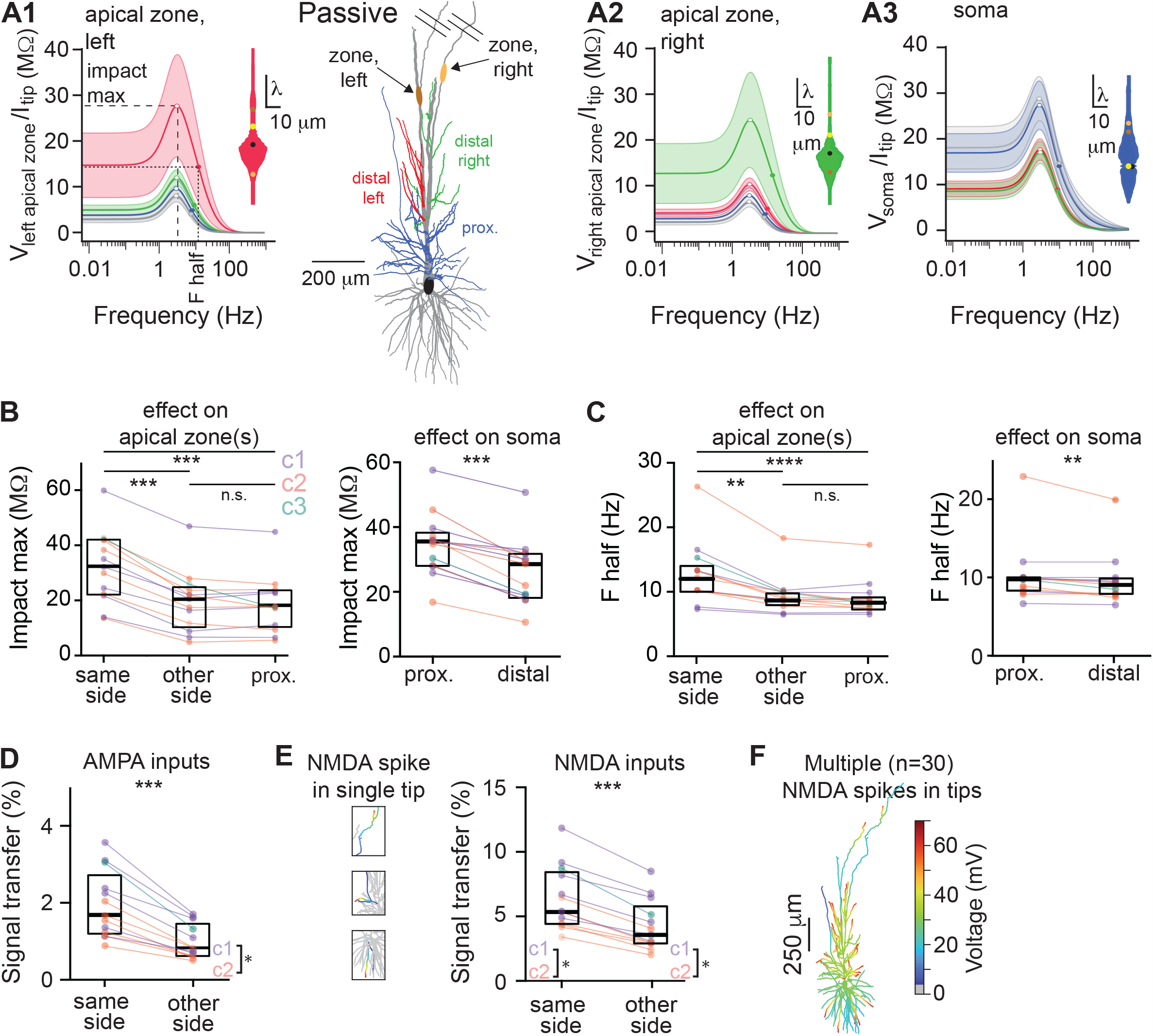
Preferential routing of subthreshold inputs in proximal and distal dendrites. (A1) Example morphology with oblique dendrites color coded as a function of location relative to the early apical trunk bifurcation. Oblique dendrites proximal to the bifurcation in blue, oblique dendrites distal to the bifurcation in red and green for left and right apical branches, respectively. A single “hotzone” (orange circles) is introduced on each apical trunk to allow activation of dendritic Ca^2+^ transients. (A1) Frequency dependent impact on the left apical hotzone in response to sinusoidal currents generated in different dendritic compartments. Effect of current injected at distal, left oblique dendrites is shown in red, effect of distal right obliques in green, proximal obliques in blue and basal dendrites in gray. Inset: “equivalent cable” as seen from one of the tips of the red obliques from the point of origin (ie tip, yellow bullet) towards left hotspot (dark brown bullet, upwards), soma (black bullet, downwards), and right hotspot (light brown bullet, downwards), respectively. (A2) Analogous to (A1) but the effect of injected current on the right hotzone. (A3) Analogous to (A1, A2) but the effect of injected current on the soma. (B, left) Population statistics for ‘impact max’ shown in A1-3 (Friedman paired test, p < 0.001, Dunn’s post-hoc). Color code of individual data points match cell cluster identity (see Fig.3). (B, right) Analogous to (B, left) but the impact of electrical signals on the soma (Wilcoxon, p < 0.001). (C, left) Population statistics for ‘F half’ in A1-3 (Friedman paired test with Dunn’s post-hoc test, p < 0.01). (C, right) Analogous to (C, left) but for the soma (Wilcoxon, p < 0.001). (D, E) Signal transfer of synaptic AMPA events (D) and NMDA spikes (E) within or between groups of oblique dendrites (Wilcoxon, p < 0.001). (F) Activation of NMDA spikes simultaneously in n=30 different tips still shows severe attenuation towards apical trunk and soma.

**Supplementary Fig. 5.**
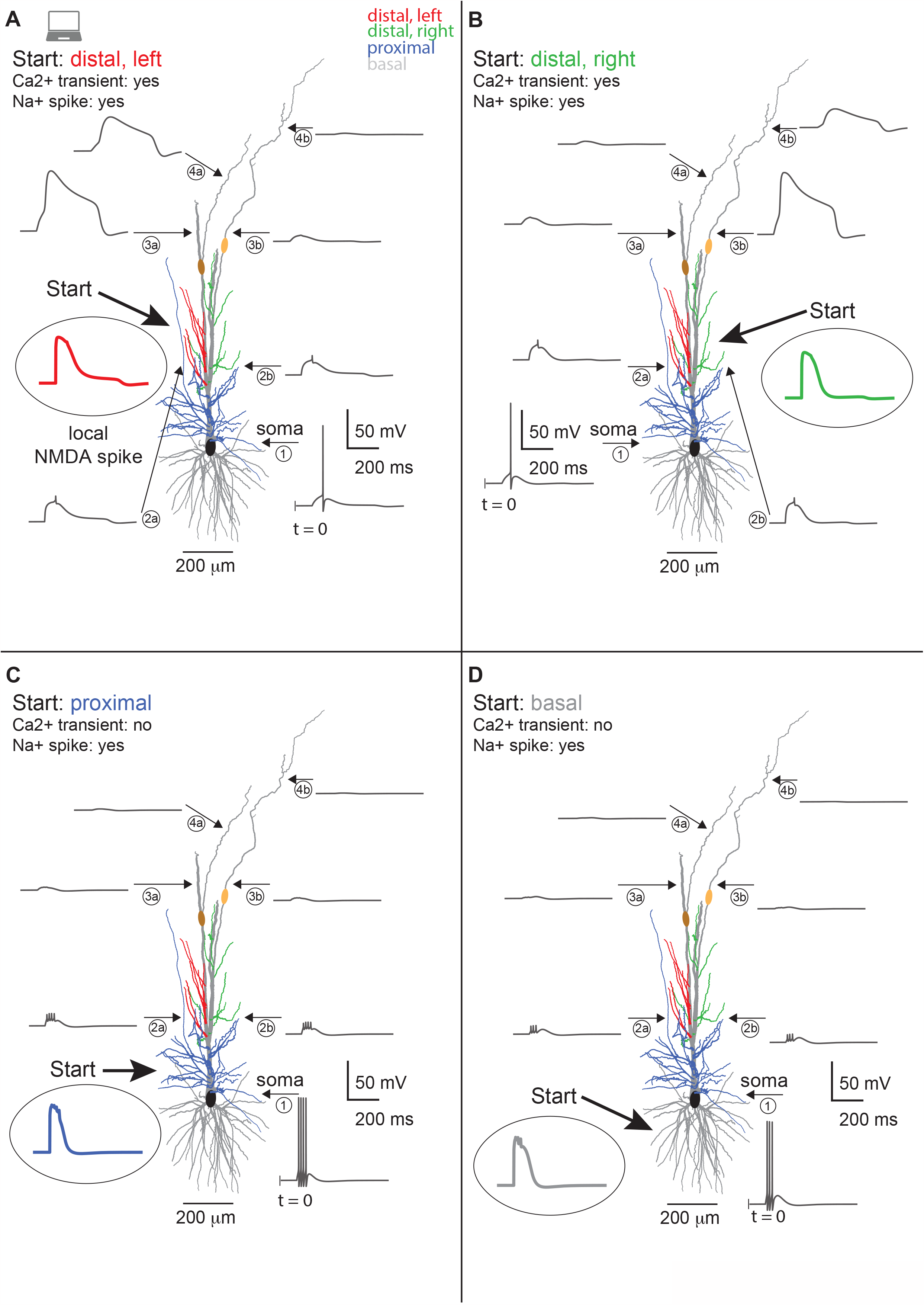
Dendritic I_h_, local NMDA spikes and Ca2+ hotspots shape non-linear output spiking. Example model (same as main Figs.4-6) with voltage traces of the simulation to illustrate the response to 7 NMDA spikes evoked in 4 different dendritic compartments: A: distal, left oblique dendrites, B) distal right oblique dendrites, C) proximal oblique dendrites and D) basal dendrites. Note that NMDA spikes in proximal obliques trigger somatic Na+ spikes, but fail to trigger apical Ca2+ transients in the hotzones. Vice versa, NMDA spikes in distal oblique dendrites trigger Ca2+ transients in the hotzones, albeit only on their respective side of origin. NMDA spikes in distal dendrites additionally trigger a somatic Na+ spike.

**Excel spreadsheet S1: Data overview. Individual datapoints are listed for each figure.**

**Suppl.Table 1:** Spine parameters for incorporating dendritic spines into human CA1 pyramidal neuron cable models, using the *F*-factor (*F = (Area*_*dendrite*_ *+ Area*_*spine*_*)/Area*_*dendrite*_).

| **Location** | **Density per 1 $\mu\text{m}$ dendritic length** | **Spine area ($\mu\text{m}^2$)** | **Distance threshold ($\mu\text{m}$)** |
| --- | --- | --- | --- |
| **Basal tree** | 2.43 | 2.12 | 60 |
| **Apical trunk** | 3.7 | 2.53 | 140 |
| **Apical collaterals** | 2.47 | 2.21 | 50 |

F was computed individually, per dendritic branch. We set $F = 1$ for all dendrites that are closer to the cell body (less than distance threshold) since spines were absent in those dendrites. Data was taken from (76).

**Suppl.Table 2:**
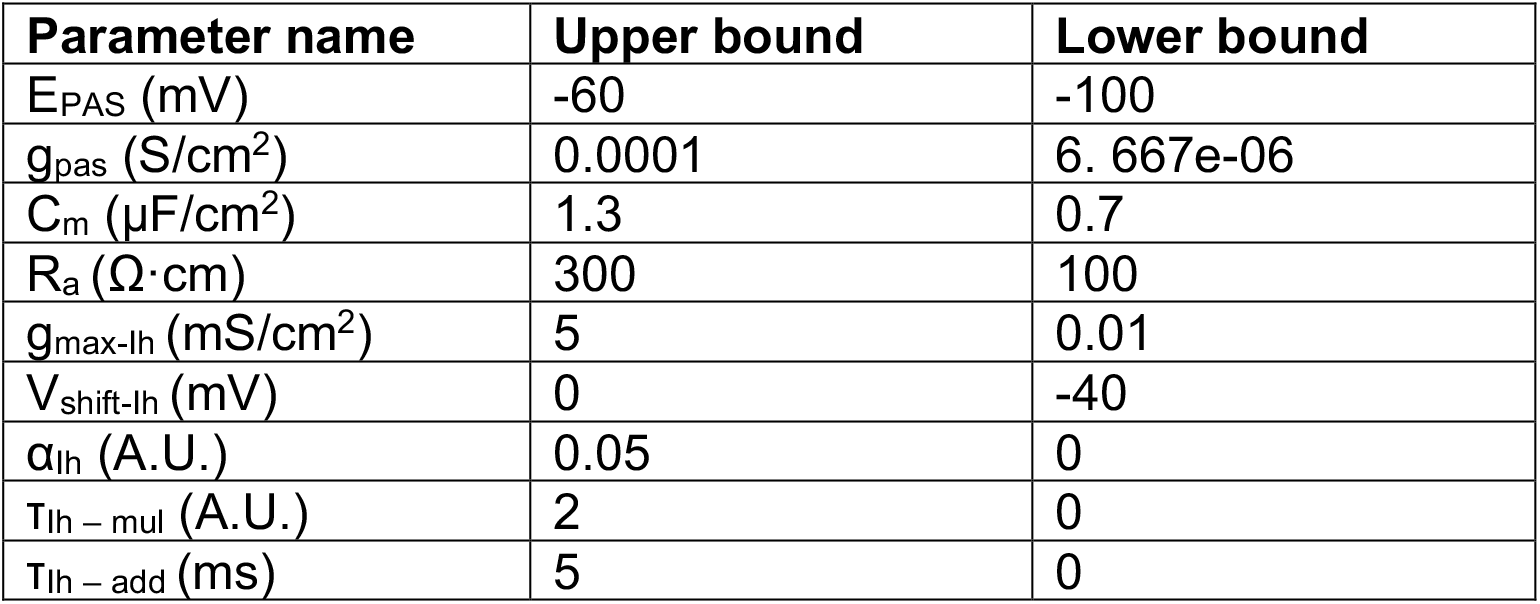
Boundaries imposed on the parameters to be optimized. Explanation on I_h_ related parameter can be found in Methods.

| **Parameter name** | **Upper bound** | **Lower bound** |
| --- | --- | --- |
| $E_{PAS}$ (mV) | -60 | -100 |
| $g_{pas}$ (S/cm<sup>2</sup>) | 0.0001 | 6. 667e-06 |
| $C_m$ (μF/cm<sup>2</sup>) | 1.3 | 0.7 |
| $R_a$ (Ω·cm) | 300 | 100 |
| $g_{max-Ih}$ (mS/cm<sup>2</sup>) | 5 | 0.01 |
| $V_{shift-Ih}$ (mV) | 0 | -40 |
| $\alpha_{Ih}$ (A.U.) | 0.05 | 0 |
| $T_{Ih - mul}$ (A.U.) | 2 | 0 |
| $T_{Ih - add}$ (ms) | 5 | 0 |

**Suppl.Table 3:** Model parameters.

| Model # | Cluster ID (Fig.3) | C<sub>m</sub> (μF/cm<sup>2</sup>) | R<sub>m</sub> (Ω·cm<sup>2</sup>) | R<sub>a</sub> (Ω·cm) | e<sub>pas</sub> (mV) | g<sub>lh_max</sub> (mS/cm<sup>2</sup>) | V<sub>shift</sub> (mV) | T<sub>add</sub> (ms) | T<sub>mul</sub> (A.U) | α<sub>lh</sub> (A.U) |
| --- | --- | --- | --- | --- | --- | --- | --- | --- | --- | --- |
| 1 | C2 | 0.7 | 139125.3 | 300 | -83.29 | 0.083 | -32.16 | 3.26 | 2 | 5.17E-10 |
| 2 | C2 | 0.7 | 149994.6 | 299.8 | -99.65 | 0.091 | -30.29 | 4.75 | 1.89 | 2.18E-07 |
| 3 | C2 | 0.7 | 149999.8 | 300 | -88.38 | 0.013 | -29.33 | 2.29 | 2 | 4.78E-07 |
| 4 | C1 | 0.7 | 149996.5 | 299.2 | -79.26 | 0.068 | -33.79 | 4.83 | 1.99 | 5.94E-07 |
| 5 | C2 | 0.7 | 149998.2 | 298.5 | -80.43 | 0.183 | -28.24 | 2.26 | 1.98 | 2.77E-13 |
| 6 | C1 | 0.7 | 107235.2 | 298.6 | -82.45 | 0.017 | -39.23 | 4.62 | 1.36 | 4.28E-07 |
| 7 | C2 | 0.71 | 26362.3 | 300 | -74.97 | 0.069 | -29.66 | 4.04 | 2 | 2.56E-14 |
| 8 | C2 | 0.7 | 90242.1 | 292.1 | -87.39 | 0.118 | -17.59 | 4.08 | 1.71 | 3.91E-10 |
| 9 | C1 | 0.77 | 80667.6 | 283.1 | -75.8 | 0.208 | -5.12 | 3.68 | 1.99 | 0 |
| 10 | C1 | 0.7 | 149969.4 | 146.2 | -88.84 | 0.072 | -23.39 | 4.82 | 1.99 | 1.23E-10 |
| 11 | C3 | 0.97 | 78278.9 | 101.7 | -82.22 | 0.289 | -11.41 | 2.58 | 1.27 | 1.24E-12 |
| 12 | C1 | 0.72 | 71380.7 | 117.1 | -79.52 | 0.106 | -21.39 | 2.94 | 2 | 0.00E+00 |
| 13 | C1 | 0.7 | 75250.2 | 201 | -79.19 | 0.136 | -4.58 | 1.82 | 1.96 | 1.45E-06 |
| 14 | C3 | 0.71 | 45320.9 | 292.9 | -76.59 | 0.05 | -30.54 | 4.79 | 1.94 | 2.89E-09 |
| 15 | C3 | 1.26 | 17857.9 | 116.5 | -75.09 | 0.29 | -17.67 | 4.86 | 1.65 | 1.9E-010 |
| 16 | C3 | 1.04 | 39510.7 | 281.54 | -76.24 | 0.01 | -36.62 | 4.98 | 2 | 1.43E-07 |

Results from fitting morphological and biophysical properties of n = 13 hCA1 pyramidal neurons. C<sub>m</sub>, R<sub>m</sub> and R<sub>a</sub> are the specific membrane capacitance, membrane resistance and axial resistance, respectively. The parameter “e<sub>pas</sub>” denotes the passive membrane battery. I<sub>h</sub> parameters, g<sub>lh\_max</sub>, V<sub>shift</sub>, T<sub>add</sub>, T<sub>mul</sub>, α<sub>lh</sub>, are explained in detail in the Methods section. Briefly, these parameters represent maximal I<sub>h</sub> somatic conductance, voltage junction potential shift, linear scale for the time constant of I<sub>h</sub> and the linear slope for the I<sub>h</sub> distribution such that g<sub>lh\_max</sub>(dendrite) = g<sub>lh\_max</sub>(soma) + distance<sub>from soma</sub> \* α<sub>lh</sub>.

